# The Evolutionary Structure of Acoustic Learnability: A Deep Learning Approach to Neotropical Birdsong

**DOI:** 10.64898/2026.02.04.703899

**Authors:** Camilo Andrés Cortés-Parra, Héctor J. Hortúa, Juan Camilo Ríos-Orjuela

## Abstract

Passive Acoustic Monitoring offered a scalable solution for biodiversity assessment in the Neotropics, although classifying hundreds of sympatric species in complex soundscapes remained challenging. We developed a deep learning framework for large-scale avian classification by training convolutional neural networks on recordings from 667 Neotropical bird species across northern South America. Efficient-NetV2L performed best, achieving 94.48% accuracy, 94.30% macro F1-score, and 0.998 macro ROC-AUC. After training, we conducted a post hoc analysis of species-level F1-scores using phylogenetically informed models (PGLS and PGLMM) to test morphological, ecological, and geographic predictors under phylogenetic control. Broad traits explained little interspecific variation: the best-supported PGLS accounted for approximately 2.2% of the variance, whereas the null model received the strongest support among PGLMM candidates. Geographic range size showed the most consistent negative association with F1-score, while morphological and ecological predictors added little explanatory power. Performance nevertheless showed weak but significant phylogenetic signal, and frequent confusions tended to involve more closely related species. Grad-CAM and Monte Carlo Dropout provided complementary descriptions of saliency and predictive uncertainty. Overall, deep learning performed effectively for regional biodiversity monitoring and provided a comparative framework for assessing how much of the variation in acoustic classification performance could be explained by broad biological predictors.

## 1 Introduction

Biodiversity monitoring in megadiverse regions, such as the Neotropics, faces a structural crisis due to systematic biases that limit the detection of temporal changes in species diversity (Abbad et al., 2025). This challenge is further exacerbated by rapid habitat loss and climate change, which are driving profound biodiversity changes across tropical ecosystems (Abbott & Le Maitre, 2010). Traditional inventory methods are logistically prohibitive and insufficient to capture biodiversity dynamics at the required spatial and temporal scales (Sugai et al., 2019). Passive Acoustic Monitoring (PAM) has emerged as a transformative solution, enabling the collection of bioacoustic data on an unprecedented scale (Ross et al., 2023). However, this technology changes the challenge from data collection to data processing, generating massive volumes of information that require automated and efficient analysis (Sugai et al., 2019).

Convolutional Neural Networks (CNNs) have become a dominant approach for acoustic signal classification, commonly approached as an image classification task using spectrograms (Kahl et al., 2021; Mac Aodha et al., 2018; Sugai et al., 2019). Nevertheless, their application in the acoustically complex and taxonomically diverse ecosystems of the Neotropics presents formidable challenges (Abbad et al., 2025). The high richness of sympatric species, interspecific acoustic convergence, and intraspecific variability (e.g., dialects) test the ability of models to learn discriminative acoustic signatures (De Kort et al., 2002; Memet et al., 2022). Despite progress in the field, a gap remains in establishing a robust benchmark for large-scale classification in this context (Sugai et al., 2019). Furthermore, bridging the gap between computational performance and biological interpretation is urgent: understanding which species are computationally “cryptic” or “distinct” offers critical insights into the structure of acoustic communities and the limitations of automated monitoring (Rivera et al., 2023; Ross et al., 2023).

Beyond the engineering challenge, birdsong reflects the outcome of multiple evolutionary and ecological forces (C. Catchpole & Slater, 1995; Derryberry et al., 2018). Competing hypotheses propose distinct drivers of acoustic diversity, ranging from deterministic constraints to stochastic or culturally mediated processes (Podos & Warren, 2007; Wilkins et al., 2013). The Morphological Constraint Hypothesis (MCH) posits that anatomical features, such as body size and beak morphology, limit the acoustic space a species can occupy, especially in terms of frequency and temporal modulation (García & Tubaro, 2018; Hay et al., 2024). This view implies that song characteristics are constrained by physical properties of the sound-producing apparatus (Podos, 1997), and thus should correlate with morphological divergence. In contrast, the Acoustic Adaptation Hypothesis (AAH) argues that song evolution is shaped by the structure of the environment, which selectively favors signal properties that maximize transmission efficiency under different acoustic conditions (Brumm & Naguib, 2009; Morton, 1975). For example, lower frequencies may be favored in closed habitats to reduce reverberation and scattering, potentially leading to convergence among ecologically similar but unrelated species (Slabbekoorn & Smith, 2002; Wiley & Richards, 1978). A third perspective highlights the role of cultural evolution: in oscine songbirds, song is learned rather than genetically encoded, allowing intraspecific variation (e.g., dialects) to emerge via social transmission and innovation (Podos, 1997; Podos & Warren, 2007; Rivera et al., 2023). This can introduce additional culturally mediated variation into song structure and may weaken its correspondence with morphology or environment.

These hypotheses are not mutually exclusive. In fact, they reflect different axes of constraint (anatomical, environmental, and historical) that may operate simultaneously and interact in complex ways (Derryberry et al., 2018; Wilkins et al., 2013). Importantly, they make divergent predictions about the predictability of acoustic signals. Under MCH or AAH, the acoustic structure should be tightly linked to measurable traits or contexts, making species more learnable for biological and artificial classifiers (García & Tubaro, 2018; Hay et al., 2024; Rivera et al., 2023). In contrast, if socially transmitted or population-specific vocal variation contributes substantially to song diversity, classification performance may be less readily predicted from coarse morphological or environmental descriptors (C. K. Catchpole & Slater, 2008; Podos & Warren, 2007; Wilkins et al., 2013). This opens a key opportunity: if model performance reflects the distinctiveness of a species’ acoustic signature, then it may also carry information about the evolutionary and ecological forces shaping that distinctiveness.

In this context, classification performance (e.g., F1-score) can be interpreted as an emergent property of the interaction between song structure and background acoustic diversity (Krause et al., 1993; Memet et al., 2022; Ross et al., 2023). Species with acoustically unique songs should be easier to distinguish, while those with convergent or variable songs should pose greater challenges to the model (Rivera et al., 2023; Wilkins et al., 2013). Notably, these patterns can be compared across species in a phylogenetically informed framework to test whether acoustic “learnability” aligns with morphological divergence, habitat, or range-related proxies of cultural complexity (García & Tubaro, 2018; Hay et al., 2024). This opens a key opportunity: if model performance reflects species-level acoustic distinctiveness within the sampled assemblage, it can be used as a comparative response to evaluate whether broad biological predictors account for variation in that distinctiveness. In this sense, classification performance is treated as a comparative proxy rather than as a direct measurement of the evolutionary processes that generated song diversity.

In this study, we focus on three well-grounded hypotheses that span this conceptual space. First, we test the Morphological Constraint Hypothesis, which proposes that species with distinct morphologies, particularly larger body mass or longer beaks, are constrained to occupy narrower or lower-frequency acoustic spaces due to biomechanical limitations of the syrinx and vocal tract (Derryberry et al., 2018; García & Tubaro, 2018; Podos & Warren, 2007). This hypothesis predicts that such species will produce more predictable and distinguishable songs, making them easier to classify (Rivera et al., 2023; Wilkins et al., 2013). Second, we examine the Acoustic Adaptation Hypothesis, which posits that the structure of the habitat influences the evolution of acoustic signals: species in dense, cluttered environments (e.g., forests) may converge on low-frequency, tonal vocalizations that transmit efficiently in reverberant conditions (Brumm & Naguib, 2009; Morton, 1975; Wiley & Richards, 1978). Under this hypothesis, environmental similarity may lead to acoustic convergence and reduced classification performance (De Kort et al., 2002; Rivera et al., 2023; Wilkins et al., 2013). Third, we evaluate a Cultural Evolution Hypothesis, where variation in vocal signals arises from socially learned behaviors and population-level dialects, independent of morphological divergence (C. K. Catchpole & Slater, 2008; Podos & Warren, 2007; Rivera et al., 2023; Slabbekoorn & Smith, 2002). Here, we use geographic range size as a proxy for potential cultural complexity and dialectal fragmentation (Podos & Warren, 2007; Wilkins et al., 2013), predicting that species with larger ranges may exhibit higher intraspecific variability and thus be harder to classify (C. K. Catchpole & Slater, 2008; Rivera et al., 2023; Wilkins et al., 2013).

To evaluate these hypotheses, we developed a deep learning framework trained on a large and taxonomically diverse dataset of birdsong from 667 species across northern South America, encompassing both passerines and key non-passerine orders. With this, we address two persistent gaps in the field: the absence of robust performance benchmarks for acoustic classification in species-rich tropical assemblages and the lack of integrative studies that link model performance to macroevolutionary theory (Abbad et al., 2025; Ross et al., 2023; Wilkins et al., 2013). Rather than treating classification accuracy solely as an endpoint, we use species-level classification performance as an assemblage-conditioned comparative proxy for acoustic distinctiveness (Brumm & Naguib, 2009; C. K. Catchpole & Slater, 2008; García & Tubaro, 2018; Morton, 1975; Podos, 1997; Podos & Warren, 2007). By integrating these outputs with standardized trait data and phylogenetic information, we ask whether broad morphological, ecological, and geographic descriptors account for interspecific variation in computational learnability. Finally, using Grad-CAM and Monte Carlo Dropout (Gal & Ghahramani, 2016; Rivera et al., 2023; Selvaraju et al., 2017), we characterize selected saliency patterns and predictive uncertainty. In doing so, we evaluate deep learning not only as a classification tool but also as a comparative framework for investigating biological structure in species-level classification performance.

## 2 Methods

### 2.1 Data Acquisition and Dataset Construction

The audio recordings were obtained from Xeno-Canto (Xeno-canto Foundation for Nature Sounds, 2005–2026), an open-access repository of avian vocalizations. To ensure high signal fidelity, we strictly filtered the data to include only recordings classified as “A” (high quality). A specific dataset was constructed for this study, comprising recordings of 667 bird species distributed in northern South America, spanning both passerine and non-passerine orders. The taxonomic classification and nomenclature were aligned with the “Guía Ilustrada de la Avifauna Colombiana” (Ayerbe-Quiñones, 2022) to ensure regional biological precision.

The final dataset, after preprocessing, consisted of 215,188 spectrograms. This was divided in a stratified manner to ensure proportional representation of each species in the three subsets: training (70%), validation (15%), and testing (15%). To prevent data leakage, the split was performed at the recording level (source file) rather than the segment level. All spectrogram segments derived from a single audio file were assigned exclusively to one subset, ensuring that the model was never evaluated on segments from recordings seen during training.

### 2.2 Audio Preprocessing and Feature Engineering

Following a robust methodology for bioacoustic analysis (Baowaly et al., 2024), a standardized preprocessing pipeline was applied.

#### Segmentation and Mel Spectrogram Generation

Each audio file was segmented into non-overlapping fragments of 5 seconds (stride equal to the window length). Segments shorter than this duration were zero-padded to maintain consistent temporal dimensions. To avoid bias from excessively long recordings, a maximum of 24 segments per original file were extracted.

Each audio segment was transformed into a Mel spectrogram using a sampling rate of 32 kHz. We configured the transformation with a Fast Fourier Transform (FFT) window of 2048 samples and 128 Mel bands. A hop length of 619 samples was selected to align the temporal resolution with the target width of 256 pixels. The analysis was restricted to a frequency range of 500-12,500 Hz.

#### Normalization and Image Conversion

Amplitude was converted to the decibel scale referencing the peak amplitude of each individual segment (ref=max). Subsequently, a Min-Max normalization was applied to scale the spectrograms to a [0-255] integer range, saving them as single-channel JPEG images with a final dimension of 128 *×* 256 pixels (height *×* width).

#### Data Augmentation

To improve model generalization, data augmentation was implemented. In the spectral domain, random time masks (up to 3 masks of 10 spectrogram time bins) and frequency masks (up to 2 masks of 40 bins) were applied. Visual transformations (brightness, Gaussian blur) were applied with a probability of 80% (fig. 1) (Baowaly et al., 2024).

**Figure 1:**
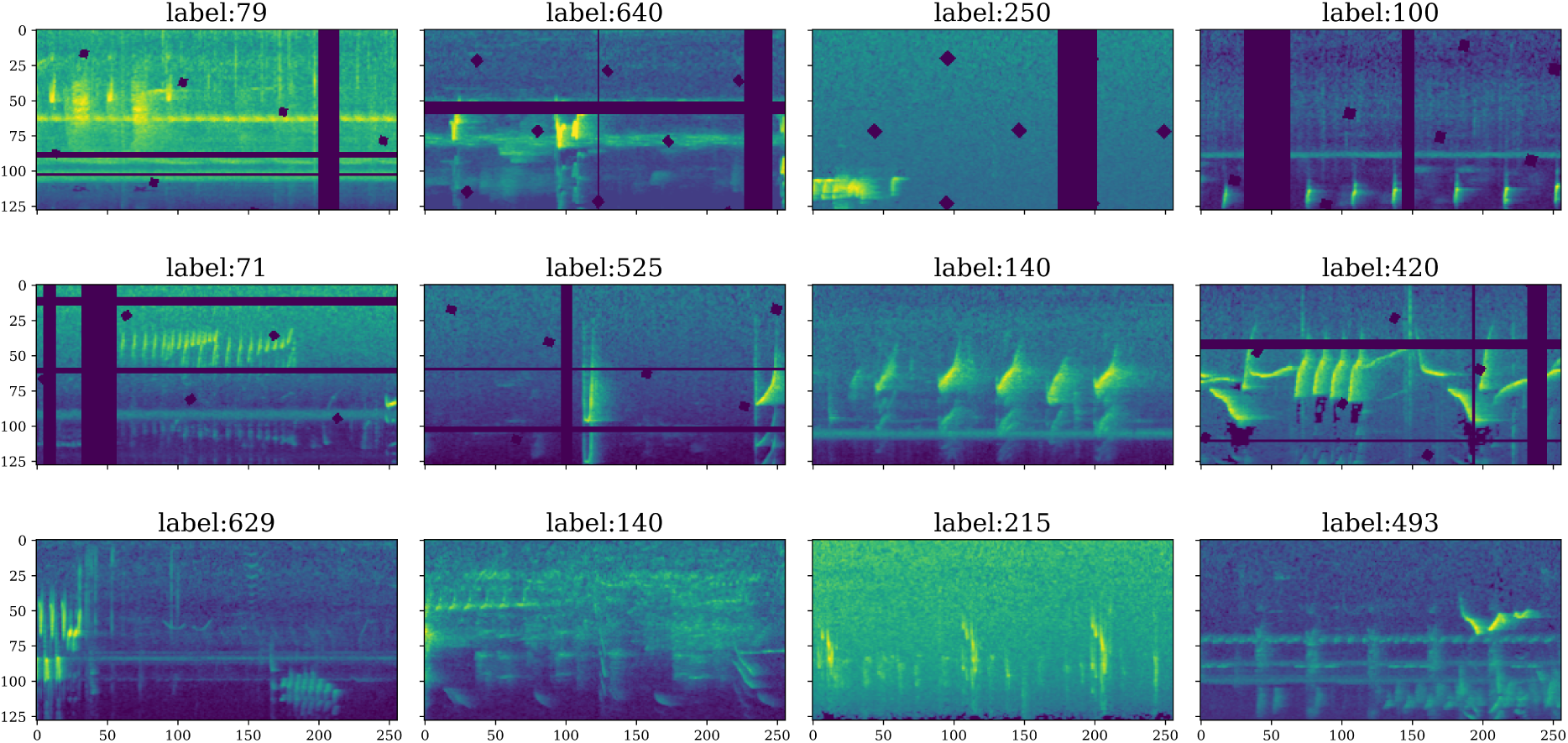
Temporal segmentation and masking generate diverse training inputs from single audio files. By slicing original recordings into 5-second non-overlapping fragments and applying spectral masks, the dataset variability is increased to improve model generalization.

### 2.3 Model Development and Training

#### Transfer Learning Architectures

Four deep CNN architectures, pretrained on ImageNet, were employed: ResNet152V2 (He et al., 2016b), MobileNetV3Large (Howard et al., 2019), EfficientNetV2L (Tan & Le, 2021), and EfficientNetB7 (Tan & Le, 2019).

#### Input Adaptation and Architecture Modifications

Unlike standard approaches that resize inputs to square dimensions, we configured the models to accept single-channel spectrograms with a shape of 128 *×* 256 pixels (height *×* width). To make these inputs compatible with the ImageNet-pretrained architectures (which expect 3-channel RGB inputs), a projection layer was integrated at the beginning of the network using a *Lambda* function to replicate the single channel into a 3-channel tensor dynamically.

The base models were instantiated with global average pooling (pooling=“avg”) to reduce dimensionality. On top of this feature extractor, a simplified classification head was added, consisting of a *Dropout* layer followed by a single dense output layer with 667 units. The models were trained with a dropout rate of 0.2. In the implementation used to generate the archived EfficientNetV2L evaluation reported here, the classification head was reconstructed with a dropout rate of 0.3 and the Dropout layer remained active during inference. Consequently, the primary EfficientNetV2L test predictions and species-level F1-scores correspond to an archived single stochastic forward pass rather than deterministic inference. Predictive variability was characterized separately through the Monte Carlo Dropout analysis described below.

#### Fine-tuning Strategy

We employed a deep fine-tuning strategy. The top 200 layers of each base architecture were unfrozen and made trainable—for MobileNetV3Large, the shallowest of the four backbones, this leaves nearly the entire network trainable (99.3% of parameters; table S2)—allowing the models to adapt their high-level feature representations specifically to the bioacoustic domain while, in the deeper backbones, preserving the low-level edge and texture detection capabilities learned from ImageNet.

#### Training and Evaluation Setup

The models were compiled using the Adam optimizer with an initial learning rate of 1*e^−^*^4^. The loss function was Categorical Cross-entropy with label smoothing (*α* = 0.1) to prevent overconfidence. Training was conducted for a maximum of 100 epochs. We implemented a set of callbacks to ensure optimal convergence: ModelCheckpoint to save the best weights based on validation accuracy, EarlyStopping with a patience of 5 epochs, and ReduceLROnPlateau to reduce the learning rate by a factor of 0.2 after 3 epochs of stagnation. All four architectures were trained using the common optimization framework described above and compared on the same held-out test partition. The archived EfficientNetV2L evaluation used for down-stream analyses retained the inference behavior described in the preceding paragraph.

### 2.4 Phylogenetic and Ecological Correlates of Performance

To evaluate associations between biological traits and model classification performance while controlling for evolutionary non-independence, we implemented phylogenetic comparative methods.

#### Phylogenetic Hypothesis and Trait Data

We sourced phylogenetic data from the global phylogeny of birds by Jetz et al. (2012), accessed via *BirdTree*. For the primary comparative analysis, we used a Maximum Clade Credibility (MCC) tree as a single fully resolved representative of the posterior tree distribution. The MCC topology is selected by maximizing the product of clade posterior probabilities and provides branch-length information from which the phylogenetic covariance matrix used in the comparative models can be constructed. We selected this approach to obtain a single resolved tree for the primary analyses while retaining posterior-supported phylogenetic structure. To evaluate sensitivity to uncertainty in this summary tree, we additionally repeated the modeling process on 10 trees randomly sampled from the posterior distribution; the qualitative interpretation of the results remained consistent across these trees.

The trees were pruned to match the intersection of species present in both our classification performance dataset (n = 667) and the trait database. This intersection yielded a final analytical set of 583 species. Therefore, it is important to note that while the global deep learning benchmark and predictive metrics are reported for the complete dataset of 667 species, the macroevolutionary comparative analyses (PGLS and PGLMM) are strictly restricted to this subset of 583 species that possess complete phylogenetic and functional trait data.

Morphological, ecological, and biogeographical traits were retrieved from the AVONET database (Tobias et al., 2022). Continuous morphological variables (body mass, culmen, tarsus, tail length, and hand-wing index) and range size were log-transformed to mitigate skewness. Categorical ecological variables (habitat, trophic level, trophic niche, and primary lifestyle) were encoded as dummy variables. To address multicollinearity, we calculated pairwise correlations, removing variables with a coefficient *>* 0.75 and prioritizing those with higher variance.

#### Phylogenetic Signal Analysis

We selected the species-level F1-score as the response variable. To quantify the phylogenetic conservation of classification performance, we estimated Pagel’s *λ* (Pagel, 1999) via maximum likelihood. The *λ* parameter ranges from 0 (phylogenetic independence) to 1 (evolution under Brownian motion). Statistical significance was assessed using a likelihood ratio test against a null hypothesis of *λ* = 0.

#### Comparative Modeling Framework (PGLS and PGLMM)

We employed two modeling approaches to identify significant predictors of the F1-score:

1. *Phylogenetic Generalized Least Squares (PGLS):* Given that F1-scores are bounded within [0, 1], we applied a logit transformation with a boundary correction (*ɛ* = 0.001) to satisfy normality assumptions:

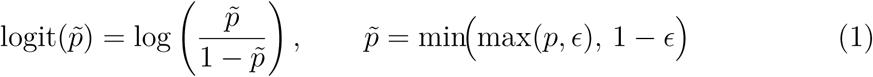 We fitted PGLS models with simultaneous estimation of *λ*, evaluating five candidate structures: Null (intercept only), Morphological, Ecological, Geographic, and a Full model.
2. *Phylogenetic Generalized Linear Mixed Models (PGLMM):* To model the F1-score on its original scale, we fitted Beta regression models incorporating a phylogenetic covariance matrix derived from the shared branch lengths. Because the Beta distribution has support on the open interval (0, 1), F1-scores exactly equal to 1 (26 species) were clipped to [10*^−^*^6^, 1 *−* 10*^−^*^6^] prior to fitting, mirroring the boundary correction applied in the PGLS.

Model selection was based on the Akaike Information Criterion (AIC). In the archived analysis, a Jacobian correction was applied to the PGLS log-likelihood to account for the logit transformation of the response. Because the PGLS and Beta-regression PGLMM nevertheless differ in response treatment and model formulation, we interpret model selection within each framework and do not use the absolute AIC difference between PGLS and PGLMM as evidence that one framework is intrinsically superior to the other.

### 2.5 Phylogenetic Confusion Analysis

Beyond overall performance, we investigated whether frequently confused species tended to be phylogenetically closer than randomly selected species pairs. Phylogenetic distance was calculated as the sum of branch lengths (in millions of years) connecting two species in the MCC tree. High-confusion pairs were defined as each species’ most frequent confusion partner in the test-set confusion matrix (i.e., one pair per species), so that each focal species contributed equally regardless of sample size. The distribution of distances among these observed confusion pairs was compared with a reference distribution generated from 1,000 randomly selected species pairs using a one-sided Mann–Whitney U test. Because this random reference does not preserve the focal-species structure of the observed confusion set, we interpret the analysis as exploratory evidence of phylogenetic structure in misclassification rather than as a generative null model of the confusion process. For the complementary within-family enrichment analysis, the expected proportion of same-family species pairs under random pairing was calculated from the observed family-size distribution (9.9%) and compared descriptively with the observed proportion of within-family misclassifications (21.1%).

The family-aggregated confusion matrix and the full MC Dropout predictive-probability matrix are provided as Supplementary Material (figs. S2 and S4).

### 2.6 Interpretability and Uncertainty Quantification

#### Gradient-weighted Class Activation Mapping (Grad-CAM)

To examine which spectro-temporal regions were associated with selected predictions, we employed Grad-CAM (Selvaraju et al., 2017). Activation maps were generated over the last convolutional layer and interpreted as illustrative saliency patterns rather than as causal evidence of the acoustic features determining model performance.

#### Uncertainty with Monte Carlo Dropout

Predictive uncertainty was characterized using Monte Carlo Dropout (Gal & Ghahramani, 2016; Hortúa et al., 2020; Seoh, 2020). The trained EfficientNetV2L weights were evaluated with the classification-head dropout rate set to 0.3, and 2,000 stochastic forward passes were performed for each sample to obtain a distribution of predictive probabilities. These distributions were used for the per-segment uncertainty analyses, species-level profiles, and corpus-level predictive-probability matrices (figs. S4 and S5). Because the inference dropout rate (0.3) differed from the rate used during model fitting (0.2), we treat these outputs as a post hoc MC Dropout-based characterization of predictive variability rather than as formally calibrated posterior probabilities.

### 2.7 Computational Platform

All experiments were conducted on the Google Colab platform, using NVIDIA A100 GPUs. Model development was carried out with the TensorFlow 2.x framework, optimizing the data pipeline with the tf.data.AUTOTUNE utility to efficiently parallelize preprocessing.

### 2.8 Evaluation Metrics and Averaging Strategy

Given the highly multiclass nature of the dataset (667 species) and the inherent class imbalance typical of crowdsourced bioacoustic data, relying solely on global accuracy can be statistically misleading. Therefore, the primary evaluation metric utilized in this study is the Macro-averaged F1-score. Threshold-independent discrimination was additionally assessed through the area under the ROC curve, computed with a one-vs-rest strategy and macro averaging, and through the area under the precision–recall curve (average precision), also macro-averaged.

To enable direct comparison with BirdNET v2.4 (Kahl et al., 2021), we additionally evaluated both models on a random recording-level sample of 11,695 recordings drawn from the held-out test partition. Each recording was processed through the same pipeline described above—segmented into 5-second fragments and converted to Mel spectrograms— over which our model produced segment-level predictions; a single recording-level prediction was then obtained by averaging the softmax probabilities across all segments of the recording and taking the argmax. Because BirdNET operates on 3-second audio windows, its window-level scores were aggregated into a single recording-level prediction by taking the maximum (top-1). Species labels were matched to the taxonomy of BirdNET by scientific name, and the comparison was restricted to the 664 study species covered by BirdNET, so that both models discriminated within the same label space. Results of this comparative evaluation are reported in the Discussion and Supplementary (table S1, fig. S6).

Biologically, Macro-F1 is the most appropriate formulation because it computes the F1-score independently for each class and then takes the unweighted mean. This ensures that rare or less frequently recorded species contribute equally to the final performance metric as highly abundant species. In the subsequent comparative analyses, each species is the analytical unit, making an unweighted species-level summary preferable to metrics dominated by the most frequently recorded taxa.

## 3 Results

### 3.1 Comparative Performance of the Evaluated Architectures

We evaluated four Convolutional Neural Network (CNN) architectures using transfer learning and compared their archived performance on the same held-out test set (table 1). EfficientNetV2L achieved the highest performance, with an overall accuracy of **94.48%** and macro-averaged precision, recall, and F1-score of **94.56%**, **94.24%**, and **94.30%**, respectively, on the test set of 32,279 segments spanning 667 species. It was followed by MobileNetV3Large (macro F1 = 92.61%), EfficientNetB7 (91.17%), and ResNet152V2 (85.81%). All four architectures also showed high threshold-independent discrimination, with macro-averaged ROC-AUC values exceeding 0.99. Training and validation curves showed no marked divergence during optimization.

**Table 1:** EfficientNetV2L attains the highest classification performance among the four evaluated architectures. The four architectures were compared on the same held-out test set (32,279 segments, 667 species). Accuracy is computed globally, whereas precision, recall, F1-score, ROC-AUC, and PR-AUC are macro-averaged across species. Architectures are ordered by macro F1-score.

| Architecture | Accuracy (%) | Macro-Prec. (%) | Macro-Rec. (%) | Macro-F1 (%) | ROC-AUC | PR-AUC |
| --- | --- | --- | --- | --- | --- | --- |
| EfficientNetV2L | 94.48 | 94.56 | 94.24 | 94.30 | 0.998 | 0.972 |
| MobileNetV3Large | 92.61 | 93.17 | 92.30 | 92.61 | 0.998 | 0.957 |
| EfficientNetB7 | 90.80 | 91.36 | 91.34 | 91.17 | 0.997 | 0.951 |
| ResNet152V2 | 86.11 | 86.45 | 85.65 | 85.81 | 0.993 | 0.903 |

A detailed engineering characterization of the evaluated architectures—including parameter counts, computational cost (FLOPs), and measured inference latency—is provided in Supplementary (table S2, fig. S1).

### 3.2 Phylogenetic Signal and Correlates of Classification Performance

The phylogenetic comparative analysis revealed significant phylogenetic signal in the classification performance (Pagel’s *λ* = 0.159, *p <* 0.001). Because *λ* is a scaling parameter of the phylogenetic covariance—ranging from 0 (phylogenetic independence) to 1 (evolution under Brownian motion)—rather than a partition of variance, this value, significantly greater than zero yet far below one, indicates a weak but statistically detectable phylogenetic signal: closely related species exhibit somewhat more similar learnability than expected under phylogenetic independence, though far less than expected under pure Brownian evolution.

#### Model Selection and Explanatory Power

Model selection was interpreted separately within the two comparative frameworks. Among the PGLS candidates, the Geographic model received the strongest support (AIC = −2397.76). Within the Beta-regression PGLMM candidate set, the intercept-only model received the strongest support (AIC = −2331.80), with no predictor set improving on its own null model. Because the two frameworks differ in response treatment and model formulation, their absolute AIC values are reported descriptively but are not used to rank PGLS against PGLMM.

Overall, the explanatory power of the tested biological variables was limited. The best-supported predictor model explained only a small fraction of the total variation 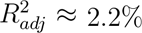, indicating that the broad morphological, ecological, and biogeographical predictors evaluated here account for little of the interspecific variation in acoustic learnability.

#### Significant Predictors

Across specifications, geographic range size showed the most consistent association with performance (*p <* 0.001 in the best-fitting Geographic PGLS): more widely distributed species were harder to classify. This association was corroborated at the coefficient level within the Beta-regression framework (geographic PGLMM: *β* = *−*0.176, *p <* 0.001), even though no PGLMM predictor set improved the AIC over its own null model (table 2); we therefore read the PGLMM fits as coefficient-level corroboration rather than model-selection support. The full model (fig. 2a) corroborated the range-size effect and surfaced two further, weaker signals: of its 26 predictors, geographic range size (*β* = *−*0.179, logit scale), the vertivore trophic niche (*β* = 1.550), and centroid latitude (*β* = 0.007) reached significance, whereas morphological variables retained no predictive power once phylogeny and the other predictors were accounted for. Sampling effort (*support*) was not treated as a biological driver but included only as a methodological covariate to absorb data-volume bias; as expected, better-represented species scored somewhat higher. Given the very low overall explanatory power, we interpret every association other than range size as a suggestive trend rather than a robust driver of learnability. For context, raw per-group averages of classification performance across habitat, habitat density, trophic level, and migratory behavior—computed without phylogenetic control—are reported in Supplementary (tables S3–S6).

**Figure 2:**
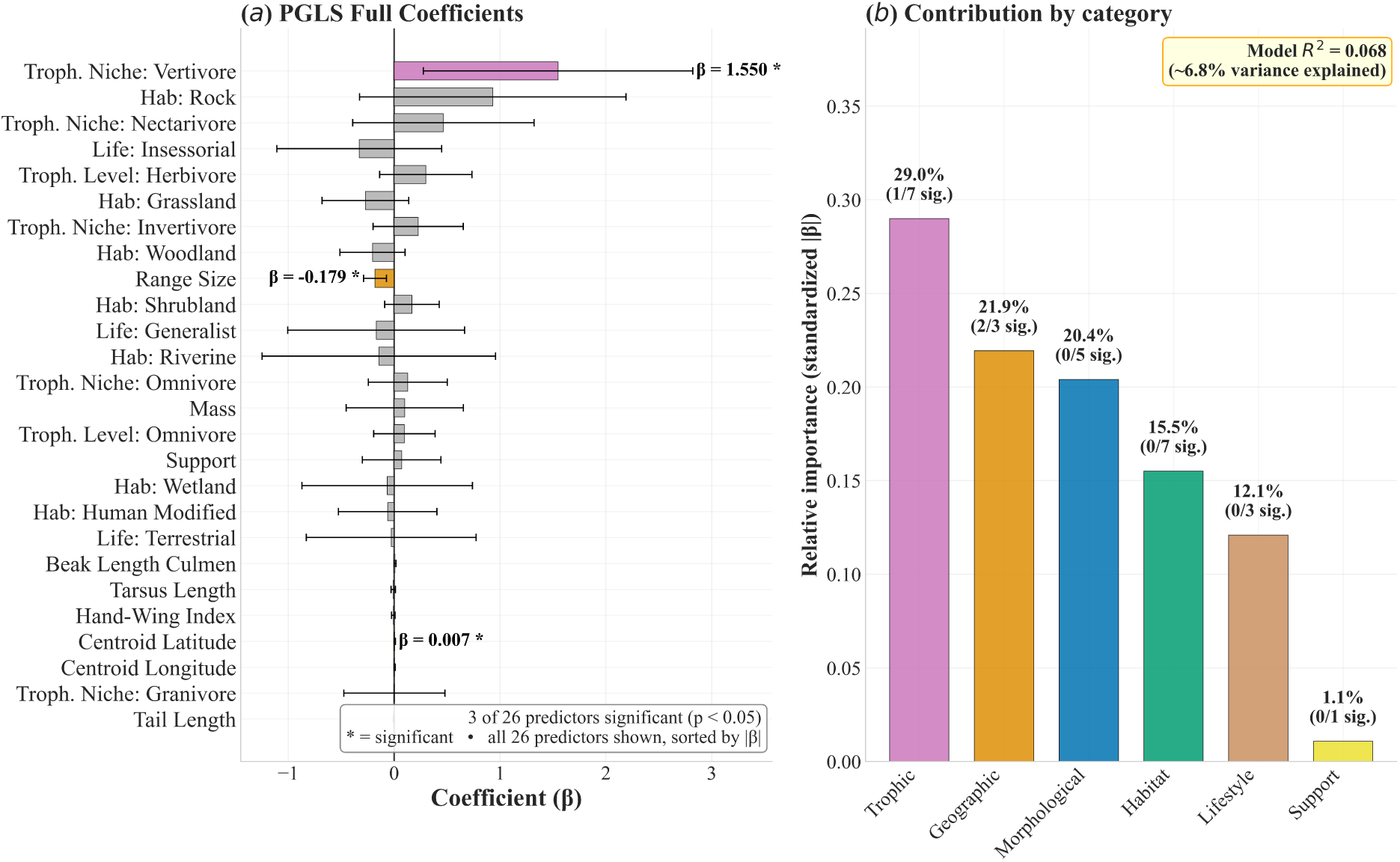
Geographic range size shows the most consistent negative association with classification performance, whereas morphological traits show little predictive contribution. (a) Raw (unstandardized) coefficients of the full PGLS model, on the logit scale, show that more widely distributed species tend to be harder to classify (*β* = *−*0.179), while body mass and beak traits are not significant; this effect was corroborated at the coefficient level in the geographic Beta-regression PGLMM (*β* = *−*0.176, *p <* 0.001). (b) Standardized effect sizes, partitioned by predictor category, indicate that geographic and trophic terms have larger standardized coefficients than the morphological terms in this model, although overall explanatory power remains low.

**Table 2:** Model selection within the PGLS and Beta-regression PGLMM frameworks. AIC values are interpreted within each modeling framework rather than as a direct ranking of PGLS against PGLMM. The Geographic model received the strongest support among PGLS candidates. Within the PGLMM candidate set, the intercept-only model received the strongest support; only this best-supported PGLMM is shown.

| Model Type | Predictor Set | AIC | ΔAIC | Key Signal |
| --- | --- | --- | --- | --- |
| PGLS | Geographic | -2397.76 | 0.00 | Range Size ($p < 0.001$) |
| PGLS | Null (Intercept) | -2388.66 | 9.10 | – |
| PGLS | Morphological | -2381.95 | 15.81 | – |
| PGLS | Full Model | -2377.57 | 20.19 | Range Size, Vertivore, Latitude |
| PGLS | Ecological | -2373.17 | 24.60 | – |
| PGLMM | Null (Intercept) | -2331.80 | – | Best-supported PGLMM |

$\Delta$ AIC values for the PGLS rows are calculated relative to the best-supported PGLS model; no cross-framework $\Delta$ AIC is reported.

### 3.3 Species-Level Performance and Outliers

Performance was near-uniformly high across the 667 species: 663 species (99.4%) attained an F1-score of at least 0.80 and 614 (92.1%) reached 0.90 or above. Twenty-six species were classified without a single error, with precision, recall, and F1-score all equal to 1.00, spanning phylogenetically diverse lineages—from antbirds and ovenbirds (*Formicivora grisea*, *Epinecrophylla ornata*, *Synallaxis albigularis*) to tanagers and finches (*Diglossa gloriosissima*, *Sphenopsis melanotis*)—indicating that flawless classification was not restricted to a single taxonomic group; a further 70 species attained perfect recall. Only four species fell below an F1-score of 0.80: *Dendrexetastes rufigula* (0.68), *Lipaugus unirufus* (0.73), *Melanospiza bicolor* (0.74), and *Molothrus oryzivorus* (0.78). At the genus level, mean performance remained high and showed only a weak descriptive relationship with the number of species per genus (linear fit *r* = 0.14; fig. S3).

### 3.4 Phylogenetic Structure of Misclassifications

We investigated whether model misclassifications showed phylogenetic structure. A Mann-Whitney U test showed that species pairs frequently confused by the model have a significantly lower phylogenetic distance compared to random pairs. The observed most-frequent confusion pairs had lower phylogenetic distances than the random-pair reference distribution (mean distance: Confused = 58.34 vs. Random = 68.48 Myr; Mann–Whitney *U* = 181,839.0, *p <* 0.001). Given the structure of the random reference described in Methods, we interpret this as exploratory evidence that frequent confusions tend to involve more closely related taxa.

This pattern is also reflected in the family-aggregated confusion matrix (fig. S2), whose dominant diagonal yields a mean family-level recall of 95.6%: only a small fraction of all predictions cross family boundaries. Within-family confusions are also enriched relative to chance—21.1% of all misclassifications involve species of the same family, 2.1 times the 9.9% expected given the observed family-size distribution. Because between-family species pairs vastly outnumber within-family ones, most errors still cross family boundaries in absolute terms; the relevant phylogenetic signal is the enrichment over chance and the reduced phylogenetic distance of confused pairs, not the absolute fraction of within-family errors.

### 3.5 Explainability and Uncertainty Quantification

To characterize model behavior beyond scalar classification metrics, we examined predictive variability using Monte Carlo Dropout and visual saliency using Grad-CAM.

At the corpus scale, the full MC Dropout predictive-probability matrix (fig. S4), aggregated by family in fig. S5, showed that the mean predictive probability assigned to the true species was 0.87 and that 90.5% of the total predictive probability mass remained within the true taxonomic family. This family-level concentration was broadly consistent with the phylogenetic structure observed in the confusion analysis.

#### Uncertainty Quantification

Species-level performance computed from the MC Dropout predictive mean was similar to the archived single-pass classification results reported above: 99.0% of species (660 of 667) achieved an F1-score *≥* 0.80, compared with 663 of 667 in the archived single-pass predictions, and no species fell below the 0.50 threshold (fig. 3). The uncertainty profiles offered granular insights into how the model handles acoustically challenging species, as illustrated in fig. 4:

- **Concentrated predictive distribution (fig. 4a):** *Volatinia jacarina* exhibited a sharply concentrated predictive distribution, with the correct class selected in 100% of the stochastic forward passes.
- **Vocal mimic example (fig. 4b):** *Mimus gilvus*, a species known for vocal mimicry, was consistently assigned to the correct class. Its predictive distribution was modestly broader than that of *V. jacarina*, indicating greater predictive dispersion in this example without identifying the biological mechanism responsible for that difference.
- **Intermediate predictive dispersion (fig. 4c):** *Riparia riparia* was correctly classified, but residual predictive probability was distributed across several alternative classes rather than concentrated on a single competitor.

**Figure 3:**
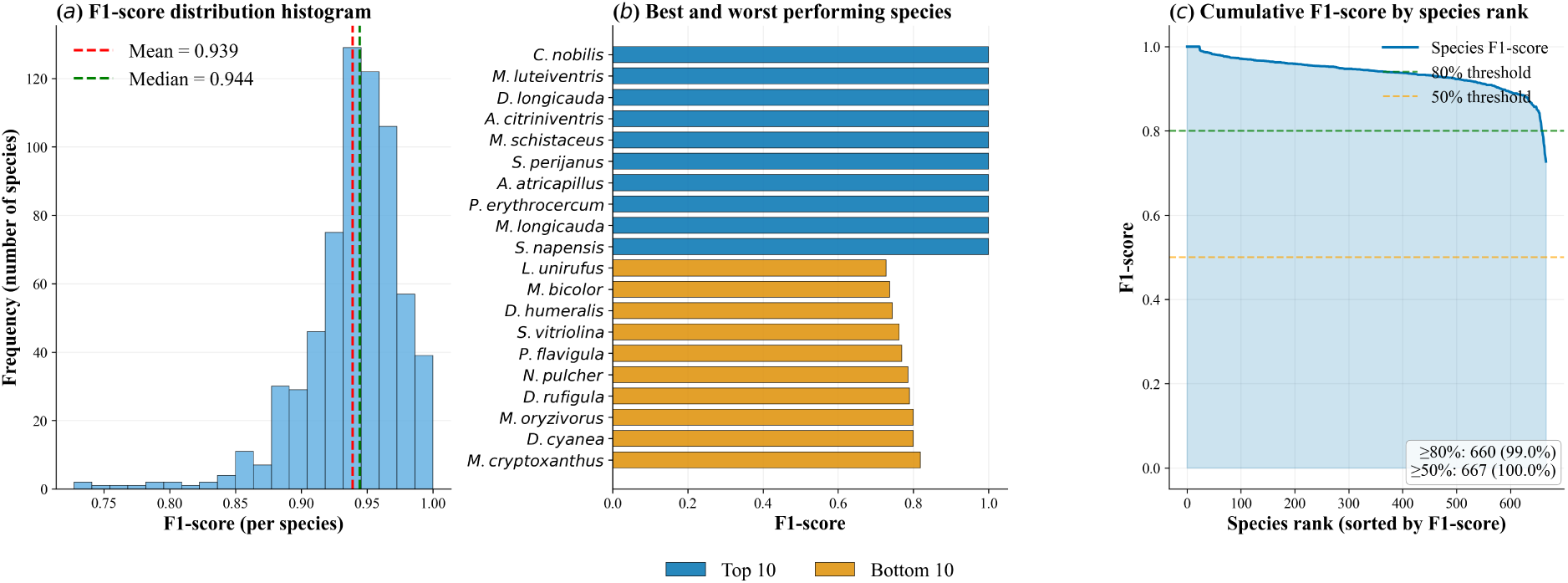
Species-level classification performance obtained from the Monte Carlo Dropout predictive mean. F1-scores in this figure are computed from the predictive probabilities averaged over 2,000 stochastic forward passes and therefore differ slightly from the archived single-pass values reported in Section 3.3 (species-level mean F1 = 0.943; 663 species *≥* 0.80). (a) The distribution is strongly skewed towards high F1-scores (mean = 0.939; median = 0.944). (b) The best- and worst-performing species span diverse taxonomic lineages. (c) 99.0% of species (*n* = 660) achieved an F1-score *≥* 0.80.

**Figure 4:**
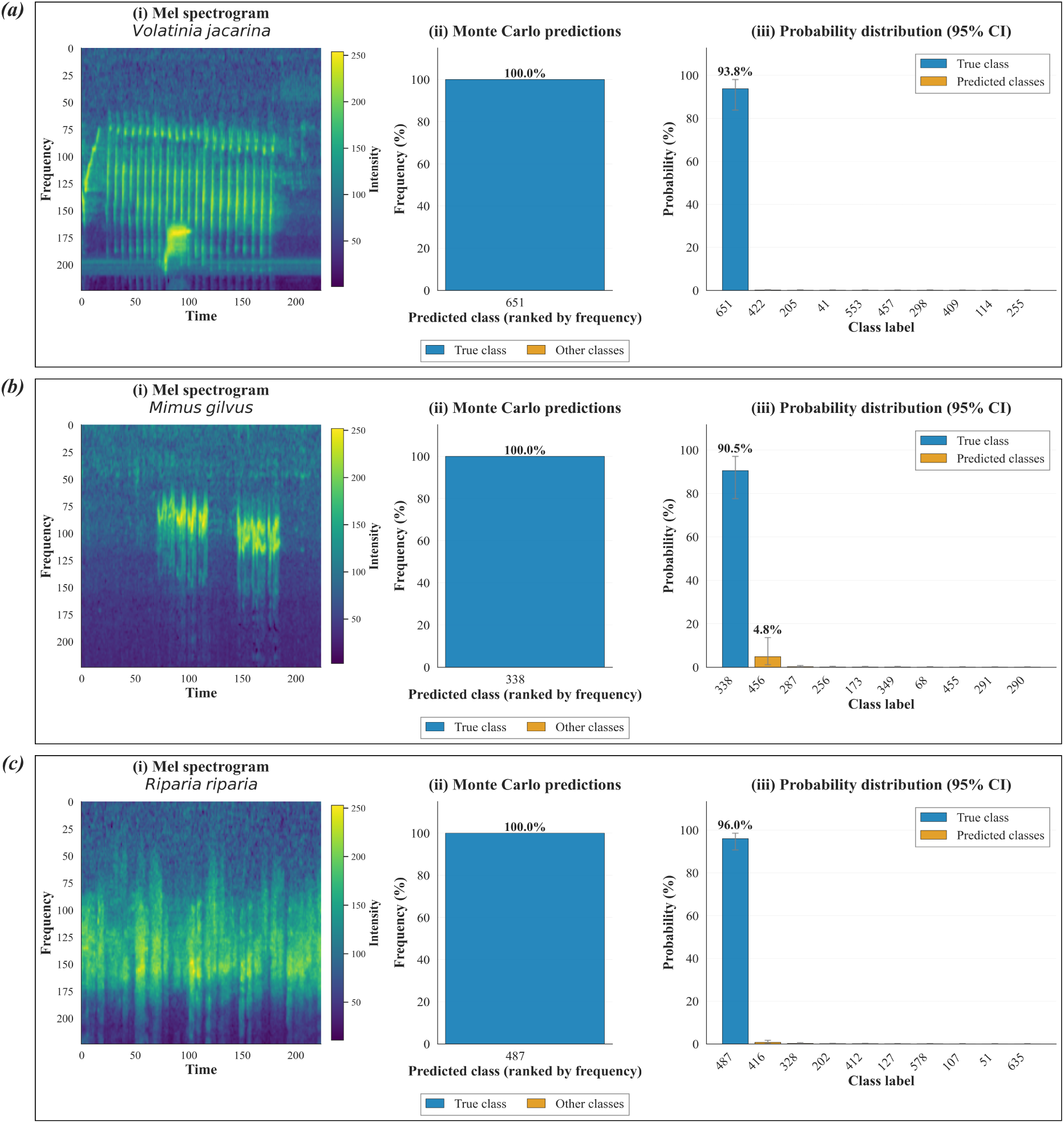
Examples of predictive distributions obtained using Monte Carlo Dropout. **(A)** *Volatinia jacarina* shows a sharply concentrated predictive distribution. **(B)** The vocal mimic *Mimus gilvus* is consistently assigned to the correct class but shows a somewhat broader predictive distribution. **(C)** *Riparia riparia* retains residual predictive probability across several alternative classes. These examples illustrate differences in predictive concentration but do not by themselves identify the biological causes of uncertainty.

#### Visual Explanations (Grad-CAM)

Grad-CAM activation maps provided illustrative examples of the spectro-temporal regions associated with individual EfficientNetV2L predictions (fig. 5). For *Acropternis orthonyx* (fig. 5a), saliency was concentrated primarily on regions overlapping the focal harmonic structure and ascending note contours. For *Anthus rubescens* (fig. 5b), activation overlapped the characteristic frequency band of the trill but was strongest over its initial notes rather than across the complete repetitive sequence. These examples are consistent with the model using spectro-temporal information contained in the focal vocalizations, although Grad-CAM alone cannot establish which acoustic features causally determine classification performance.

**Figure 5:**
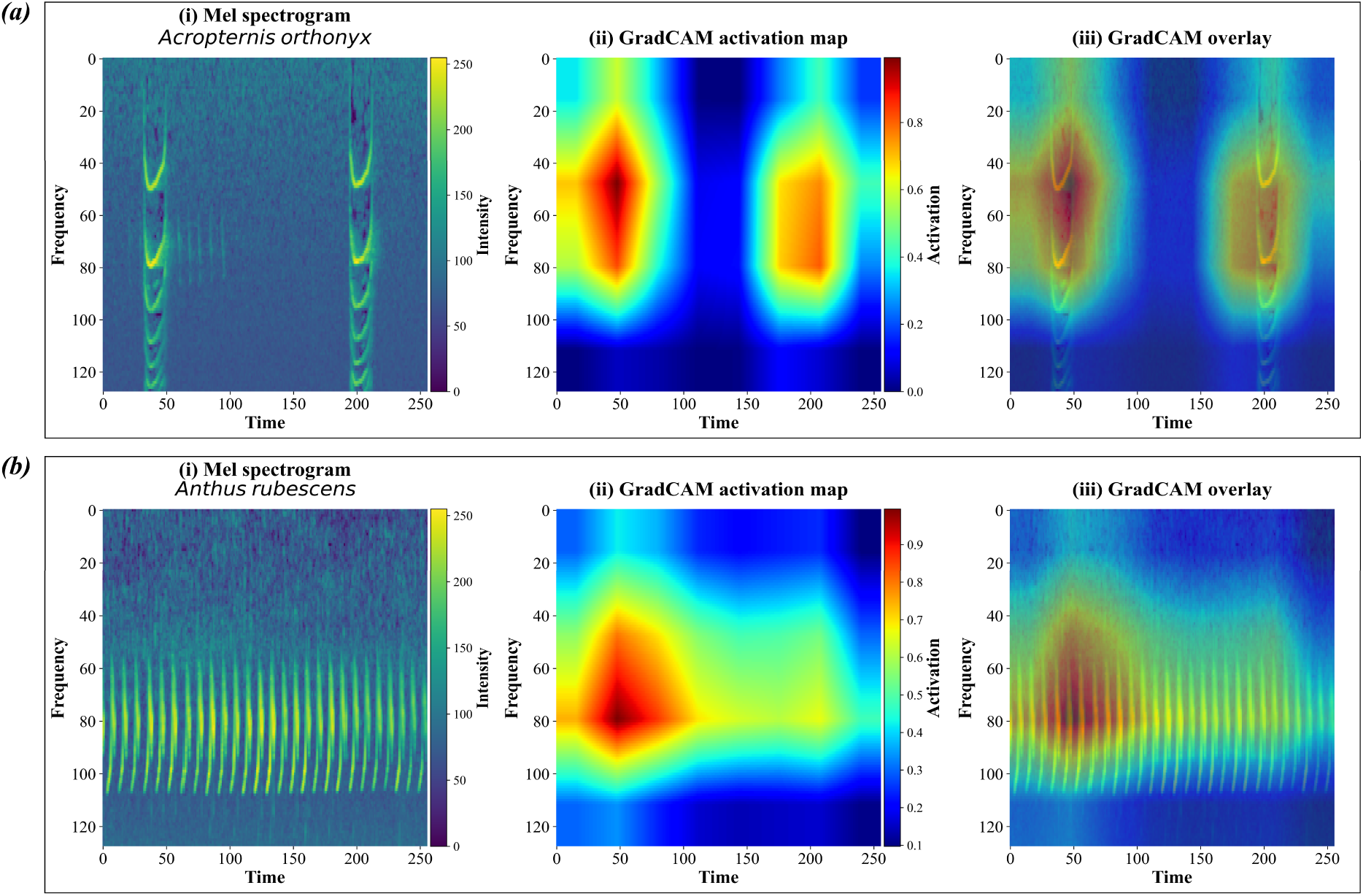
Illustrative Grad-CAM saliency maps for two EfficientNetV2L predictions. **(A)** In *Acropternis orthonyx*, saliency overlaps regions containing the focal harmonic stacks and ascending note contours. **(B)** In *Anthus rubescens*, saliency overlaps the trill’s characteristic frequency band and is strongest over the initial notes rather than across its full temporal extent. These examples are consistent with the use of spectro-temporal information from the focal vocalizations but are not interpreted as causal evidence of the features determining model performance.

## 4 Discussion

This study demonstrates the effectiveness of deep learning architectures for the large-scale classification of Neotropical birds, a task complicated by high species diversity and acoustic complexity. The performance attained by EfficientNetV2L shows that modern convolutional architectures can learn spectro-temporal representations sufficient to distinguish among hundreds of species, consistent with trends in computational bioacoustics (He et al., 2016a; Kahl et al., 2021; Mac Aodha et al., 2018). However, beyond the engineering success, our results offer a unique opportunity to explore the biological underpinnings of acoustic distinctiveness and to evaluate how well established evolutionary hypotheses align with patterns of classification performance.

### 4.1 Benchmarking in Megadiverse Regions

To contextualize the engineering performance of our model, we include BirdNET v2.4 (Kahl et al., 2021) as a technical reference point. This comparison is not intended to position our model as a replacement for BirdNET—a global pretrained bioacoustic classifier trained on an incomparably larger and more diverse corpus—but rather to situate our regionally specialized architecture within the current landscape of bioacoustic classifiers. The two tools serve fundamentally different purposes: BirdNET targets broad-scale species detection across global soundscapes, whereas our model is regionally specialized and coupled to downstream comparative and predictive-variability analyses.

When evaluated on the same random recording-level sample of 11,695 recordings drawn from the held-out test partition, restricted to the 664 study species covered by BirdNET, EfficientNetV2L achieved higher recording-level classification performance than BirdNET v2.4 in this regional benchmark (accuracy = 0.957 vs. 0.903; macro F1 = 0.952 vs. 0.900). This comparison is intended as a paired technical benchmark rather than as evidence of general superiority: BirdNET is designed for a substantially broader taxonomic and geographic scope, whereas our model is regionally specialized. Because Xeno-Canto material contributes to BirdNET’s training resources, potential overlap between its training corpus and individual recordings in this evaluation cannot be excluded; accordingly, the comparison should not be interpreted as fully independent external validation. Confidence scores also differed between models. In this benchmark, EfficientNetV2L showed stronger separation between top-1 confidence distributions for correct and incorrect predictions than BirdNET v2.4 (fig. S6): median confidence was 0.96 for correct and 0.32 for incorrect EfficientNetV2L predictions, compared with 0.99 and 0.63, respectively, for BirdNET. Top-1 confidence discriminated correct from incorrect predictions with ROC-AUC = 0.967 and PR-AUC = 0.998 for EfficientNetV2L, versus 0.881 and 0.984 for Bird-NET v2.4. These quantities characterize confidence–correctness discrimination and are not multiclass classification AUCs. Our workflow additionally incorporates Monte Carlo Dropout, Grad-CAM, and phylogenetic analyses of classification errors as complementary tools for examining model behavior.

Situating our results more broadly, Ardila-Villamizar et al. (2026) report a macro F1-score of 0.868 for CNN-based classifiers on an independent Neotropical dataset, and Cañas et al. (2025) report a ROC-AUC of 0.936 for the Middle Magdalena BirdCLEF+ evaluation. Because this ROC-AUC value was obtained under a different multi-taxon dataset and evaluation protocol, it is provided only as descriptive context and is not directly comparable with the confidence–correctness AUCs reported for the paired BirdNET benchmark.

### 4.2 Biological Correlates of Acoustic Learnability

To investigate biological correlates of classification difficulty, we evaluated predictor sets representing morphology, environment, and geographic distribution. Classification performance showed weak but significant phylogenetic signal (Pagel’s *λ* = 0.159, *p <* 0.001). Within the PGLS candidate set, the Geographic model received the strongest support (AIC = −2397.76), whereas within the Beta-regression PGLMM candidate set the intercept-only model received the strongest support. We therefore do not interpret the absolute AIC difference between the two frameworks as evidence that one framework is intrinsically superior. Geographic range size showed the most consistent negative association with performance, whereas morphological and ecological predictor sets provided little explanatory improvement. Even the best-supported predictor model accounted for only a small fraction of the interspecific variation in F1-score 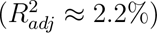.

Contrary to the expectations of the Morphological Constraint Hypothesis, our phylogenetically controlled models showed no significant association between anatomical features and classification performance. While expectations suggested that larger birds may be easier to classify, consistent with the idea that size-related constraints shape frequency and temporal patterns (García & Tubaro, 2018; Podos, 1997), no morphological trait retained consistent predictive power across model frameworks once common ancestry was controlled for. This aligns with prior studies emphasizing the importance of phylogenetic signal in acoustic structure (Derryberry et al., 2018; García & Tubaro, 2018), and indicates that the broad allometric and biomechanical proxies evaluated here provide little predictive information about classification performance. Overall, the broad morphological variables evaluated here provide little predictive information about interspecific variation in computational learnability once phylogenetic structure is considered.

Similarly, our results offered limited support for the Acoustic Adaptation Hypothesis. Although species in dense habitats might be expected to converge on low-frequency tonal signals for optimal transmission (Brumm & Naguib, 2009; Morton, 1975; Wiley & Richards, 1978), habitat structure was not a significant predictor in any of our phylogenetic models. This absence of detectable association may partly reflect insufficient ecological resolution; coarse habitat classifications from global databases fail to capture the microenvironmental complexity that shapes signal propagation (Brumm & Naguib, 2009; Wiley & Richards, 1978). Thus, this should not be interpreted as a falsification of the hypothesis, but rather highlights that coarse ecological proxies are insufficient to predict computational learnability.

Finally, we evaluated the Cultural Evolution Hypothesis, which suggests that widespread species, with greater opportunities for population fragmentation and dialect formation, may exhibit higher intra-specific variability (C. K. Catchpole & Slater, 2008; Rivera et al., 2023). The direction of the association matched this prediction: geographic range size showed a significant inverse relationship with F1-score—a pattern compatible with the “cultural trap” idea where ecological expansion challenges signal recognition (Podos & Warren, 2007)—and the coefficient remained significant in the best-fitting PGLS and was corroborated at the coefficient level in the Beta-regression PGLMM, even though no PGLMM predictor set outperformed its own null model in AIC terms. Given the sampling confounds acknowledged below, we treat this as directional support rather than direct evidence of cultural processes. The apparent correlation with trophic niche (Derryberry et al., 2018; Wilkins et al., 2013), where specialized diets seemingly influence acoustic stereotypy (Tobias et al., 2014; Wilkins et al., 2013), was weaker and specification-dependent, reaching significance only in the full PGLS; we therefore treat it as suggestive rather than robust. Range-restricted species might still present more uniform vocal profiles (C. K. Catchpole & Slater, 2008; Wilkins et al., 2013), but ultimately, these functional proxies fail to capture the vast majority of the variance in acoustic distinctiveness.

### 4.3 Limited Explanatory Power of Broad Biological Predictors

A central result of this study is the low explanatory power of the broad biological predictors evaluated here: the best-supported Geographic PGLS accounted for approximately 2.2% of the variation in species-level F1-score. This result indicates that coarse species-level descriptors of morphology, ecology, and geographic distribution are insufficient to account for most interspecific variation in computational acoustic learnability. At the same time, the weak but significant phylogenetic signal and the tendency for frequently confused species to be more closely related than the random-pair reference indicate that classification performance is not phylogenetically unstructured.

The unexplained component may encompass multiple sources of variation not represented in our comparative dataset, including fine-scale acoustic structure, population-level geographic variation, behavioral context, recording heterogeneity, annotation error, and other historical or ecological processes. Vocal learning, social complexity, sexual selection, cultural evolution, and vocal plasticity are plausible contributors in some lineages (Derryberry et al., 2018; Podos & Warren, 2007; Wilkins et al., 2013), but these mechanisms were not quantified directly in the present study and therefore remain hypotheses for future testing.

### 4.4 Phylogenetic Structure of Errors and Data Constraints

Misclassification patterns provided exploratory evidence of phylogenetic structure. The most-frequent confusion pairs were more closely related than the random-pair reference, and within-family confusions were enriched relative to chance, even though most individual errors crossed family boundaries in absolute terms. Because the random-pair reference does not preserve the focal-species structure of the observed confusion set, these results are interpreted as a tendency rather than as a definitive generative test of the confusion process. The pattern is consistent with greater similarity in classification-relevant acoustic information among close relatives, but the present analysis cannot distinguish shared ancestry, convergence, or other causes of acoustic similarity (De Kort et al., 2002; Wilkins et al., 2013).

Monte Carlo Dropout provided a complementary description of predictive variability. Residual predictive probability was disproportionately concentrated within the true taxonomic family, broadly consistent with the confusion analysis. Individual species differed in the concentration of their predictive distributions, but these differences cannot be attributed specifically to mimicry, spectral overlap, annotation uncertainty, or other biological mechanisms without targeted analyses. Predictive variability may therefore help identify cases that warrant closer biological or data-quality investigation rather than diagnose the cause of ambiguity directly.

Finally, sampling effort played a detectable but modest role: species with fewer recordings tended to score lower, a known limitation in deep learning applications to biodiversity (Mac Aodha et al., 2018), although support accounted for little of the explained variation once phylogeny and the other predictors were controlled for (fig. 2b). Improving representation of rare or range-restricted taxa nonetheless remains essential (Abbad et al., 2025; Sugai et al., 2019). Active learning or data augmentation may offer pathways forward (Kahl et al., 2021; Mac Aodha et al., 2018).

### 4.5 Limitations and Future Directions

While our findings offer new insights into the evolutionary structure of acoustic learnability, some limitations must be acknowledged. First, classification performance is not an intrinsic species trait but an assemblage-conditioned proxy: a species’ F1-score reflects the distinctiveness of its song relative to the particular set of co-occurring species sampled here, as well as recording effort—the latter mitigated by including support as a covariate. It is also partly classifier-dependent: species-level F1-scores correlated only moderately across the four evaluated architectures (r = 0.40–0.59), so our comparative analyses characterize the learnability landscape captured by the best-performing model rather than an architecture-invariant trait. This relational nature is nonetheless biologically meaningful, as acoustic niches are inherently defined relative to sympatric taxa. Second, our trait dataset, though standardized and phylogenetically comprehensive, relies on species-level averages and lacks intra-specific resolution. This constrains our ability to model processes such as sexual dimorphism, seasonal variation, or fine-scale ecological specialization, all of which are known to influence vocal signals in birds (C. K. Catchpole & Slater, 2008; Wilkins et al., 2013). Third, we did not explicitly quantify structural attributes of the vocalizations themselves, such as trill rate, frequency bandwidth, modulation patterns, or note diversity. Without these acoustic-level variables, hypotheses related to biomechanical trade-offs or acoustic niche partitioning within communities remain inaccessible (Memet et al., 2022; Podos, 1997). Incorporating automated acoustic feature extraction pipelines into future workflows could address this gap and open the door to more fine-grained comparative analyses. Fourth, our test of the cultural evolution hypothesis relies on geographic range size as a proxy for dialectal diversity. Although this proxy is grounded in empirical precedent, it does not capture within-species vocal variation directly (Podos & Warren, 2007; Wilkins et al., 2013), and it is sensitive to a methodological confound: widely distributed species tend to be recorded by more observers, with more heterogeneous equipment and acoustic environments, which could by itself reduce classification performance without invoking cultural processes. Moreover, because the test was applied across the entire assemblage—including suboscine and non-passerine lineages whose vocalizations are innate rather than learned—support for the cultural evolution hypothesis should be regarded as indirect; explicit tests restricted to vocal-learning clades would strengthen this inference. Future studies using population-level song recordings and diversity indices would allow for more explicit testing of cultural complexity and its impact on learnability. Fifth, the analysis was restricted to the 500–12,500 Hz frequency band, so species that vocalize predominantly outside this range (e.g., low-frequency tinamous and doves, or high-frequency hummingbirds) may be mischaracterized for technical rather than biological reasons.

Additionally, species-level F1-scores were estimated from finite test sets (8–75 segments per species), so sampling error may affect estimated effect sizes, phylogenetic signal, and statistical power; the comparative models do not explicitly propagate this measurement uncertainty. In addition, because the classification-head Dropout layer remained active in the archived evaluation used to generate the primary EfficientNetV2L predictions, the species-level F1-scores entering the comparative analyses correspond to a single stochastic realization of the trained model. The separate 2,000-pass MC Dropout analysis characterizes predictive variability but this inference-level variability was not propagated through the downstream phylogenetic models. Also, recording-level labels were inherited by all segments of a file, with no explicit filter for segments lacking the focal species; combined with per-segment peak normalization, this may introduce label noise from background species or low-signal fragments. Our uncertainty analysis derives from a single Dropout layer at the classification head and uses an inference dropout rate of 0.3, compared with 0.2 during model fitting. We therefore interpret these distributions as a post hoc characterization of predictive variability rather than as formally calibrated posterior uncertainty; deeper stochastic approximations or model ensembles could yield different uncertainty profiles. Finally, data imbalance and annotation quality remain persistent challenges (Abbad et al., 2025; Mac Aodha et al., 2018). Despite controlling for sample size in our analyses, rare or understudied species may be underrepresented or misannotated in the training data, limiting both performance and biological inference. Addressing these challenges will require collaborative data sharing, improved annotation standards, and possibly semi-supervised or active learning approaches tailored for biodiversity data (Mac Aodha et al., 2018; Ross et al., 2023).

Furthermore, while our dataset split strictly segregates data at the recording file level to prevent segment leakage, it is important to acknowledge the limitations inherent to crowdsourced repositories like Xeno-Canto. These databases generally lack persistent individual identifiers across different audio files. Consequently, it remains possible that multiple separate recordings of the exact same individual bird (e.g., recorded sequentially by the same observer) were inadvertently distributed across the training, validation, and test splits. This potential overlap could inflate the reported classification performance if the deep learning model is partially memorizing individual-specific acoustic signatures rather than purely generalizing at the species level. Future bioacoustic workflows should aim to utilize datasets with explicit individual metadata to perform individual-aware cross-validation, strictly isolating species-level generalization from individual recognition.

### 4.6 Biological Models from Machine Learning

This study uses deep learning as both a classification tool and a source of quantitative responses for comparative biological analysis. Species-level classification performance, predictive distributions, saliency maps, and error structure provide complementary descriptions of how the trained model behaves across taxa. These quantities should not be treated as direct measurements of evolutionary mechanisms, but they can be integrated with phylogenetic and trait data to generate and evaluate biological hypotheses. In the present study, this approach shows that broad morphological, ecological, and geographic descriptors account for little of the observed interspecific variation in computational learnability, while phylogenetic structure remains detectable. Future studies incorporating direct acoustic measurements, population-level variation, and improved sampling designs will be necessary to identify the mechanisms underlying the remaining variation.

### 4.7 Conclusion

This study shows that deep learning models, when regionally optimized, can achieve high accuracy in classifying birdsong across acoustically complex and taxonomically diverse Neotropical communities. Among the four evaluated CNN architectures, EfficientNetV2L provided the strongest performance on the regional corpus.

By employing a post hoc analytical workflow that couples species-level CNN performance metrics with phylogenetically aware comparative models, we used computational classification difficulty as an assemblage-conditioned proxy for acoustic distinctiveness. Broad morphological, ecological, and geographic variables accounted for only a small proportion of interspecific variation: geographic range size showed the most consistent negative association with performance, but the best-supported predictor model retained very low explanatory power and the null model was preferred within the PGLMM candidate set. Classification performance also showed weak but significant phylogenetic signal, and the exploratory confusion analysis indicated that frequently confused species tended to be more closely related than the random-pair reference. These results indicate that the coarse biological descriptors evaluated here are insufficient to explain most variation in computational acoustic learnability. The remaining variation may reflect fine-scale acoustic, behavioral, geographic, historical, methodological, and other unmeasured factors that require direct evaluation in future studies.

Overall, our two-stage approach supports the use of deep learning both for regional species identification and as a comparative framework for investigating biological structure in classification difficulty. Rather than replacing broad-coverage classifiers such as BirdNET, the regional model provides a complementary system in which classification performance, predictive variability, saliency, and phylogenetic error structure can be examined jointly. These analyses highlight both the strong practical performance of regional acoustic classifiers and the limited ability of coarse macroecological and morphological traits to predict interspecific variation in computational learnability.

## Ethics

This work did not require ethical approval from a human subject or animal welfare committee.

## Data Accessibility

Full data and analysis code are openly available in the GitHub repository (C. Cortés-Parra, 2026) and archived on Zenodo (C. A. Cortés-Parra, 2026).

## Acknowledgements

We are deeply grateful to the Escuela Colombiana de Ingeniería Julio Garavito for being our academic home. We also extend our heartfelt appreciation to the faculty members of the Master in Data Science program. Their mentorship, wisdom, and unwavering support throughout our studies were instrumental not only in this project but in our growth as researchers.

## Declaration of AI Use

During the preparation of this manuscript, an AI-based language tool was utilized exclusively for proofreading and to improve the grammar, clarity, and readability of the English text. The AI tool had no role in the scientific analysis, interpretation of results, or the formulation of the core ideas and conclusions presented in this work.

## Authors’ Contributions

C.C.P.: Conceptualization, Data Curation, Formal analysis, Methodology, Software, Validation, Visualization, Writing – original draft, Writing – review & editing.

H.H.: Conceptualization, Formal analysis, Supervision, Writing – review & editing.

J.C.R.O.: Conceptualization, Formal analysis, Investigation, Supervision, Writing – review & editing.

All authors gave final approval for publication and agreed to be held accountable for the work performed therein.

## Conflict of Interest Declaration

We declare we have no competing interests.

## Funding

The research and publication of this article were funded entirely by the lead author, C.C.P.

## Notes

### Competing Interest Statement

The authors have declared no competing interest.

### Summary of Updates

The manuscript has been revised and following a complete internal audit of the analyses and reporting. The model benchmark EfficientNetV2L is now identified as the best performing architecture, with updated results reported consistently throughout the manuscript. All downstream species level, phylogenetic, uncertainty, and explainability analyses now refer to this model.

