## Supplementary material for "The Evolutionary Structure of Acoustic Learnability: A Deep Learning Approach to Neotropical Birdsong"

### A Supplementary Material

The supplementary material provides the technical validation of the model, a comparative benchmarking analysis, and an ecological breakdown of acoustic learnability. We summarize the comparative performance of EfficientNetV2L and the other evaluated architectures using classification-performance and computational-cost analyses. A classification performance comparison between EfficientNetV2L, BirdNET v2.4, and related studies is provided (Table S1), along with a confidence-separation analysis illustrating the confidence-separation comparison between EfficientNetV2L and BirdNET (Fig. S6). Furthermore, descriptive summaries of classification success across ecological groupings (habitat, habitat density, trophic level, and migratory behavior) are provided for context (tables S3–S6); these raw averages do not control for phylogenetic non-independence and should be read in light of the comparative analyses presented in the main text. Finally, a comprehensive report detailing precision, recall, and F1-scores for all 667 Neotropical species is included.

#### A.1 Supplementary Figures

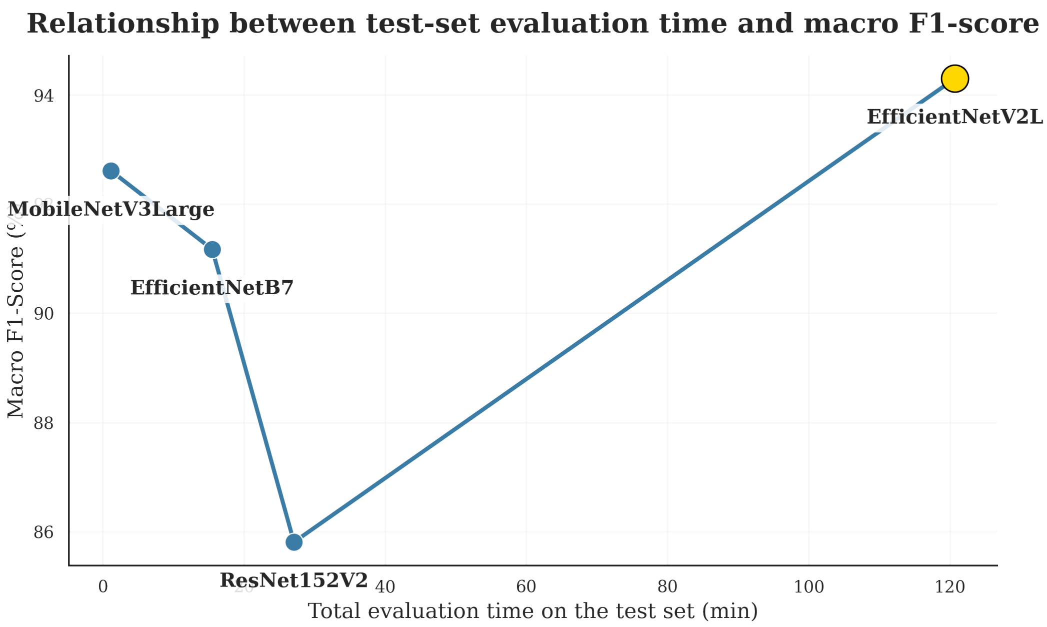


Figure S1: Trade-off between computational cost and classification performance across the four evaluated architectures. Each point relates the total wall-clock time required to evaluate the full test set (32,279 segments) to the macro F1-score achieved. EfficientNetV2L attains the highest macro F1-score at the highest computational cost, whereas MobileNetV3Large offers the fastest evaluation.

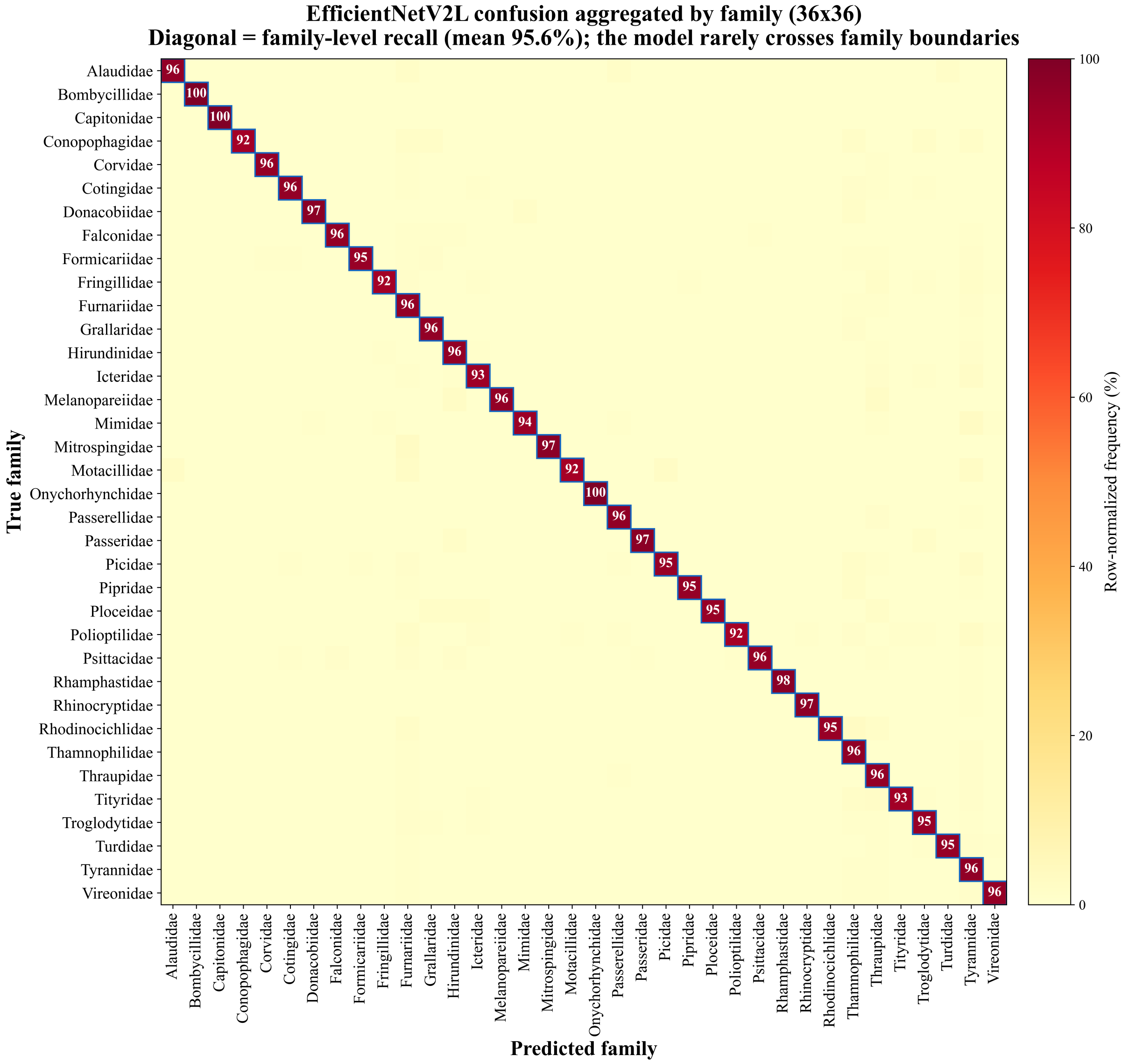


Figure S2: **Family-aggregated confusion matrix (36 × 36).** Species are collapsed into their families; the diagonal is the family-level recall (mean 95.6%), showing that the model rarely crosses family boundaries.

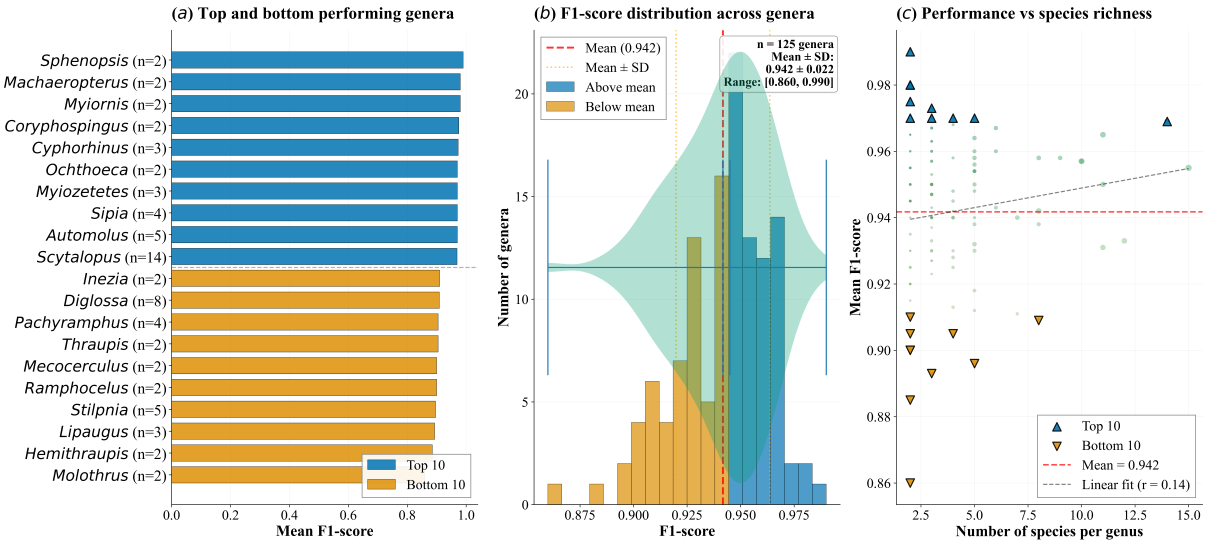


Figure S3: Descriptive genus-level classification performance and species richness. (a) Sphenopsis and Machaeropterus show the highest mean F1-scores among the displayed genera, whereas Molothrus and Hemithraupis show the lowest. (b) The distribution of mean F1-scores across 125 genera centers on a mean of 0.942 (± 0.022 SD), with a range of [0.860, 0.990]. (c) Mean F1-score shows only a weak descriptive relationship with the number of species per genus (linear fit $r = 0.14$). No inferential test is presented here, so this panel should not be interpreted as evidence of statistical independence.

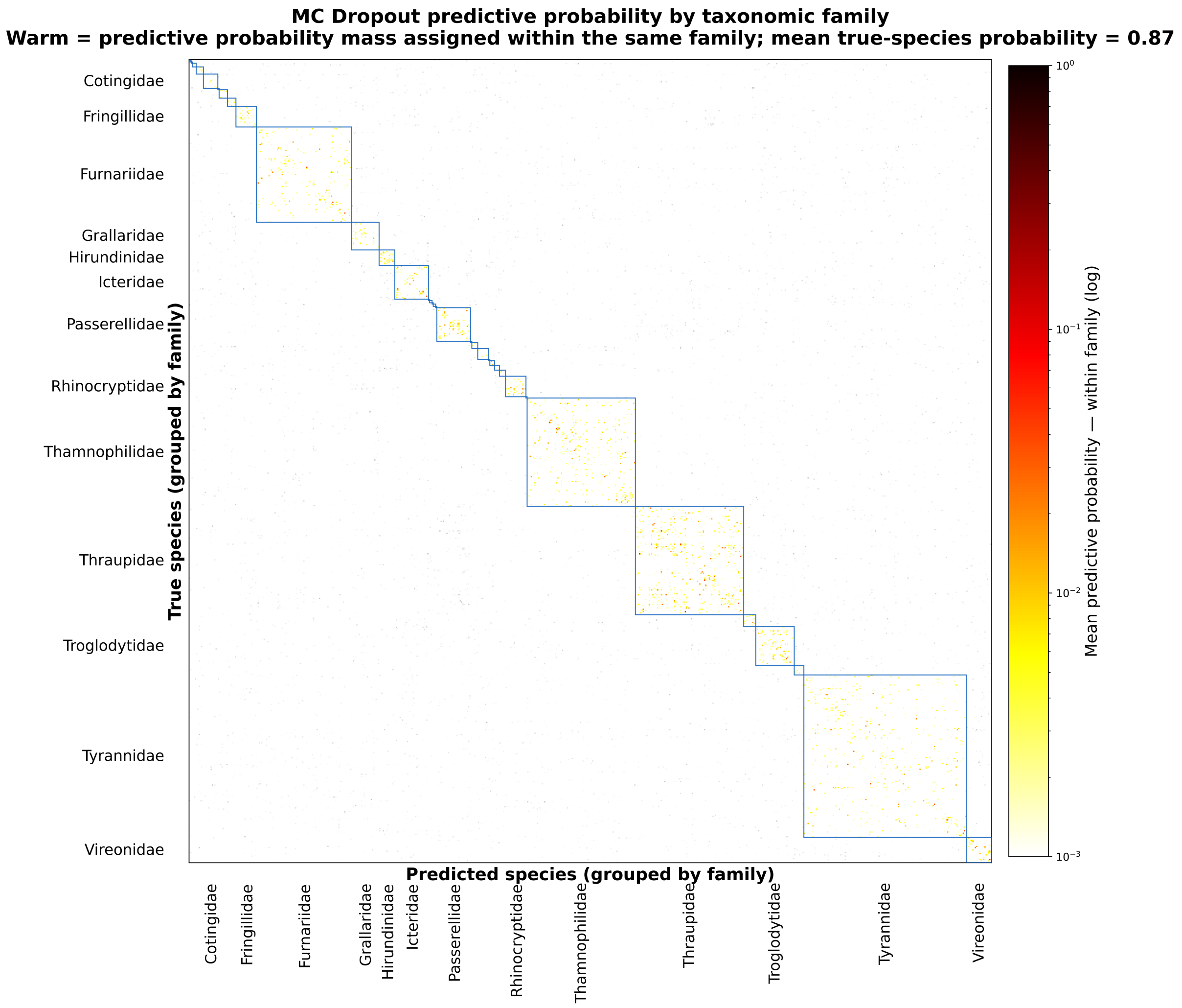


Figure S4: **MC Dropout predictive-probability matrix for EfficientNetV2L (667 species).** Warm cells show the mean predictive probability mass assigned within the same family (log scale); between-family mass appears in pale tones, and the correct diagonal is omitted from the color scale. Mean predictive probability assigned to the true species = 0.87. Predictive probability is visibly structured across taxonomic family blocks.

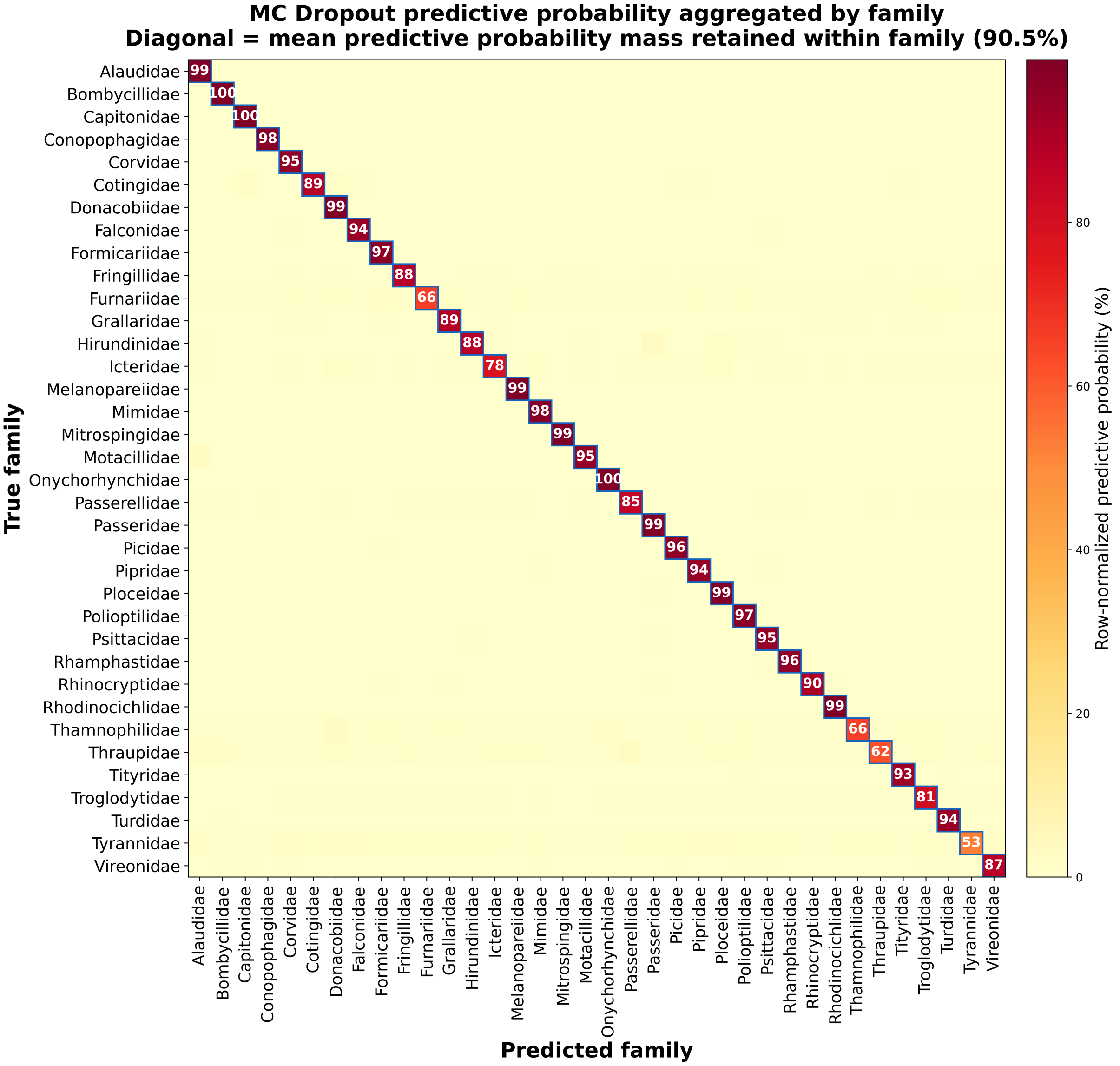


Figure S5: **Family-level aggregated predictive-probability matrix under MC Dropout.** Diagonal values represent the mean predictive probability mass retained within the true taxonomic family (90.5%). Predictive probability is disproportionately concentrated within family blocks, a pattern broadly consistent with the phylogenetic structure observed in the classification errors.

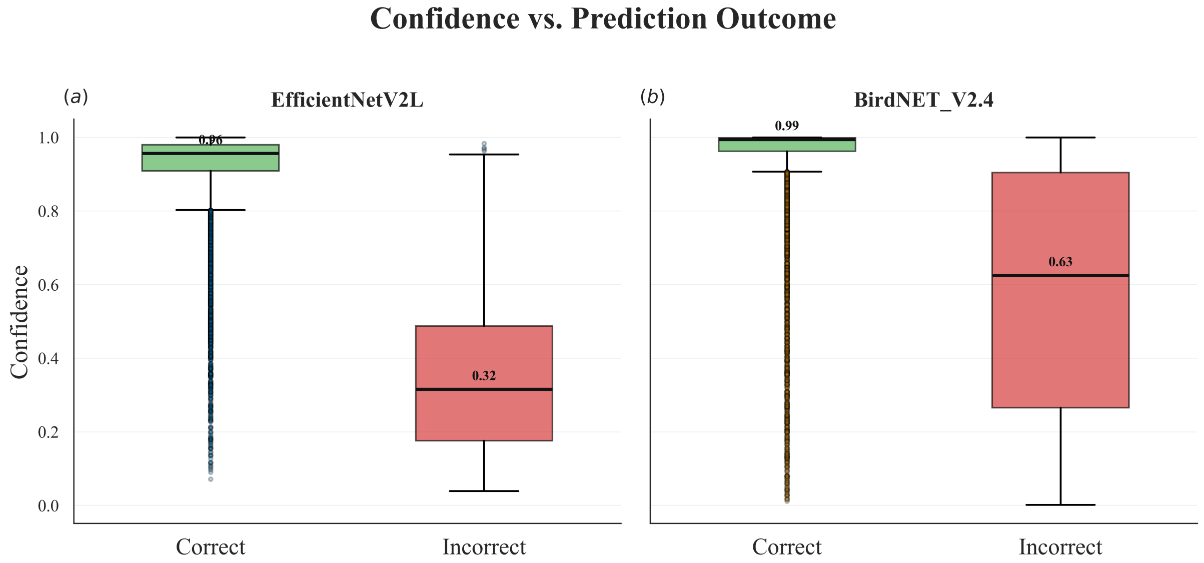


Figure S6: **Confidence distributions for correct and incorrect predictions in EfficientNetV2L and BirdNET v2.4.** Both models were evaluated on the same random recording-level sample of 11,695 recordings drawn from the held-out test partition. Top-1 confidence discriminated correct from incorrect predictions with ROC-AUC = 0.967 and PR-AUC = 0.998 for EfficientNetV2L, versus 0.881 and 0.984 for BirdNET v2.4. Median confidence was 0.96 for correct and 0.32 for incorrect EfficientNetV2L predictions, compared with 0.99 and 0.63, respectively, for BirdNET. EfficientNetV2L therefore showed stronger separation between confidence distributions for correct and incorrect predictions in this benchmark. These AUC values quantify confidence–correctness discrimination and should not be interpreted as multiclass classification ROC-AUC or PR-AUC.

#### A.2 Supplementary Tables

Table S1: **Comparative classification performance across models evaluated on Neotropical bird assemblages.** Metrics are macro-averaged unless otherwise indicated. Dashes (—) indicate metrics not reported in the original study. EfficientNetV2L and BirdNET v2.4 were evaluated on the same random recording-level sample of 11,695 recordings drawn from the held-out test partition, with both models restricted to the 664 study species covered by BirdNET; these values differ from those in Table 1, which reports segment-level performance on the full test partition (32,279 segments, 667 species). †For EfficientNetV2L and BirdNET v2.4, PR-AUC and ROC-AUC quantify the top-1 confidence as a discriminator of correct versus incorrect predictions (not macro one-vs-rest classification AUCs; see fig. S6). Ardila-Villamizar et al. (2026) report results on a separate Neotropical dataset (only macro F1-score available); Cañas et al. (2025) report a classification ROC-AUC of 0.936 on a multi-taxon competition dataset from the Middle Magdalena region, a different quantity from the confidence-discrimination AUCs marked with †. These cross-study values are provided only for descriptive context because the datasets, label spaces, aggregation procedures, and evaluation protocols differ.

| **Model / Study** | **Dataset** | **Accuracy** | **Macro F1** | **Macro Prec.** | **Macro Rec.** | **Weighted F1** | **Weighted Prec.** | **Weighted Rec.** | **PR-AUC** | **ROC-AUC** |
|---|---|---|---|---|---|---|---|---|---|---|
| EfficientNetV2L (this study) | 664 spp., N. South America | 0.957 | 0.952 | 0.957 | 0.951 | 0.957 | 0.960 | 0.957 | 0.998† | 0.967† |
| BirdNET v2.4 (this study) | 664 spp., N. South America | 0.903 | 0.900 | 0.912 | 0.899 | 0.901 | 0.909 | 0.903 | 0.984† | 0.881† |
| Ardila-Villamizar et al. (2026) | 295 spp., Neotropical (independent) | — | 0.868 | — | — | — | — | — | — | — |
| Cañas et al. (2025) | 147 spp., Middle Magdalena, Colombia | — | — | — | — | — | — | — | — | 0.936 |

Table S2: **Engineering characterization of the evaluated architectures.** Total and trainable parameters, proportion of trainable parameters, and computational cost (FLOPs for a single forward pass at the working input resolution of $128 \times 256$). The proportion of trainable parameters differs across architectures because the fine-tuning strategy unfreezes the top 200 layers of each backbone: for MobileNetV3Large, the shallowest backbone, this leaves nearly its entire network trainable, whereas the deeper backbones retain a substantial frozen ImageNet base. FLOPs were computed with the TensorFlow profiler (a multiply–accumulate is counted as two operations; equivalently 7.14, 8.05, 3.44 and 0.15 GMACs, respectively). Inference latency is reported as the mean ± standard deviation per image at batch size 1 on an NVIDIA A100 GPU; at batch size 16, throughput reaches 480, 34, 34 and 25 images/s for MobileNetV3Large, ResNet152V2, EfficientNetB7 and EfficientNetV2L, respectively. Latency and throughput were measured on pre-loaded input tensors and thus reflect pure model inference; the total wall-clock time of a full test-set evaluation (fig. S1), which additionally includes image decoding and data-pipeline overhead, is correspondingly higher. Direct runtime comparisons with published studies were not attempted because reported inference times depend strongly on hardware, batch size, preprocessing, and input dimensions; Table S2 therefore emphasizes controlled within-study comparisons under identical hardware and input conditions.

| **Architecture** | **Total params** | **Trainable params** | **Trainable (%)** | **FLOPs (G)** | **Latency (ms/img)** |
|---|---|---|---|---|---|
| ResNet152V2 | 59.70 M | 32.85 M | 55.0 | 14.29 | 57.60 ± 1.11 |
| EfficientNetV2L | 118.60 M | 55.18 M | 46.5 | 16.09 | 75.00 ± 2.37 |
| EfficientNetB7 | 65.81 M | 45.32 M | 68.9 | 6.89 | 54.97 ± 1.21 |
| MobileNetV3Large | 3.64 M | 3.61 M | 99.3 | 0.29 | 5.78 ± 0.13 |

Table S3: **Descriptive mean F1-score by habitat category.** Values are unadjusted group means and do not control for phylogenetic non-independence or other covariates.

| **Habitat Type** | **Average F1-score** |
|---|---|
| Rock | 98.00 |
| Riverine | 95.67 |
| Human Modified | 94.47 |
| Forest | 94.38 |
| Woodland | 94.25 |
| Grassland | 94.12 |
| Wetland | 94.00 |
| Shrubland | 93.89 |

Table S4: **Descriptive mean F1-score by trophic level.** Mean classification performance varied little among the three trophic categories; these unadjusted averages do not control for phylogenetic non-independence or other covariates.

| **Trophic Level** | **Average F1-score** |
|---|---|
| Carnivore | 94.64 |
| Omnivore | 93.85 |
| Herbivore | 93.68 |

Table S5: **Descriptive mean F1-score by habitat-density category.** Mean classification performance was similar across categories; these unadjusted averages do not control for phylogenetic non-independence or other covariates.

| **Habitat Density** | **Average F1-score** |
|---|---|
| Open (3) | 94.53 |
| Dense (1) | 94.50 |
| Semi-open (2) | 93.95 |

Table S6: **Descriptive mean F1-score by migratory behaviour.** Values are unadjusted group means and should not be interpreted as evidence of an effect of migratory behaviour on classification performance.

| **Migratory Behaviour** | **Average F1-score** |
|---|---|
| Migratory | 95.29 |
| Sedentary | 94.37 |
| Partial Migration | 93.03 |

#### A.3 Detailed Classification Performance per Species

The following table presents the precision, recall, and F1-score for all 667 species evaluated using the EfficientNetV2L architecture.

Table S7: Individual species metrics reveal high variability in acoustic learnability across the Neotropical dataset (EfficientNetV2L).

| species | precision | recall | f1-score | support |
|---|---|---|---|---|
| Acropternis orthonyx | 1.00 | 0.99 | 0.99 | 75 |
| Amblycercus holosericeus | 0.95 | 0.78 | 0.86 | 27 |
| Ammodramus aurifrons | 0.97 | 0.97 | 0.97 | 58 |
| Ammodramus humeralis | 0.91 | 0.93 | 0.92 | 74 |
| Ammodramus savannarum | 0.97 | 0.95 | 0.96 | 75 |
| Anabacerthia striaticollis | 0.96 | 0.93 | 0.94 | 27 |
| Anabacerthia variegaticeps | 0.93 | 0.98 | 0.96 | 44 |
| Anairetes parulus | 1.00 | 0.89 | 0.94 | 36 |
| Andigena nigrirostris | 0.91 | 0.97 | 0.94 | 63 |
| Anisognathus igniventris | 0.97 | 1.00 | 0.99 | 34 |
| Anisognathus lacrymosus | 0.86 | 0.92 | 0.89 | 26 |
| Anisognathus somptuosus | 0.90 | 0.97 | 0.94 | 39 |
| Anthus lutescens | 0.91 | 0.83 | 0.87 | 24 |
| Anthus rubescens | 0.90 | 1.00 | 0.95 | 27 |
| Ara ararauna | 0.95 | 0.93 | 0.94 | 75 |
| Arremon assimilis | 0.99 | 0.95 | 0.97 | 75 |
| Arremon atricapillus | 1.00 | 1.00 | 1.00 | 24 |
| Arremon aurantiirostris | 0.91 | 0.95 | 0.93 | 75 |
| Arremon brunneinucha | 0.94 | 0.97 | 0.95 | 75 |
| Arremon castaneiceps | 1.00 | 0.96 | 0.98 | 27 |
| Arremon taciturnus | 0.96 | 0.99 | 0.97 | 75 |
| Arremonops conirostris | 0.96 | 0.93 | 0.95 | 75 |
| Asemospiza fuliginosa | 0.93 | 0.89 | 0.91 | 46 |
| Asemospiza obscura | 0.95 | 0.95 | 0.95 | 40 |
| Asthenes flammulata | 0.87 | 0.96 | 0.91 | 27 |
| Asthenes fuliginosa | 0.92 | 0.98 | 0.95 | 45 |

| species | precision | recall | f1-score | support |
|---|---|---|---|---|
| Asthenes wyatti | 0.94 | 0.97 | 0.95 | 30 |
| Atalotriccus pilaris | 0.92 | 0.94 | 0.93 | 35 |
| Atlapetes albofrenatus | 0.96 | 1.00 | 0.98 | 26 |
| Atlapetes flaviceps | 0.93 | 0.96 | 0.95 | 27 |
| Atlapetes fuscoolivaceus | 0.97 | 0.94 | 0.95 | 32 |
| Atlapetes latinuchus | 0.97 | 0.97 | 0.97 | 75 |
| Atlapetes leucopis | 1.00 | 0.92 | 0.96 | 38 |
| Atlapetes leucopterus | 0.89 | 0.89 | 0.89 | 18 |
| Atlapetes melanocephalus | 1.00 | 1.00 | 1.00 | 35 |
| Atlapetes pallidinucha | 0.95 | 0.96 | 0.96 | 56 |
| Atlapetes schistaceus | 0.95 | 0.97 | 0.96 | 59 |
| Attila bolivianus | 0.96 | 0.90 | 0.93 | 49 |
| Attila cinnamomeus | 0.95 | 0.93 | 0.94 | 75 |
| Attila citriniventris | 1.00 | 1.00 | 1.00 | 49 |
| Attila spadiceus | 1.00 | 0.93 | 0.97 | 75 |
| Attila torridus | 0.93 | 1.00 | 0.96 | 39 |
| Aulacorhynchus albivitta | 0.97 | 0.97 | 0.97 | 75 |
| Automolus infuscatus | 0.97 | 0.99 | 0.98 | 69 |
| Automolus melanopezus | 0.96 | 0.96 | 0.96 | 27 |
| Automolus ochrolaemus | 0.99 | 0.99 | 0.99 | 75 |
| Automolus rufipileatus | 1.00 | 0.96 | 0.98 | 51 |
| Automolus subulatus | 0.93 | 0.95 | 0.94 | 44 |
| Bombycilla cedrorum | 0.92 | 1.00 | 0.96 | 45 |
| Brotogeris jugularis | 0.95 | 0.97 | 0.96 | 75 |
| Buthraupis montana | 0.97 | 0.95 | 0.96 | 37 |
| Cacicus cela | 0.95 | 0.92 | 0.93 | 75 |
| Cacicus haemorrhous | 0.88 | 0.92 | 0.90 | 75 |
| Cacicus oseryi | 1.00 | 1.00 | 1.00 | 28 |
| Cacicus solitarius | 0.96 | 0.88 | 0.92 | 51 |
| Cacicus uropygialis | 0.92 | 0.97 | 0.94 | 35 |
| Campephilus melanoleucos | 0.99 | 0.96 | 0.97 | 75 |
| Camptostoma obsoletum | 0.94 | 0.96 | 0.95 | 75 |
| Campylorhamphus procurvoides | 0.90 | 1.00 | 0.95 | 27 |
| Campylorhamphus pusillus | 0.93 | 0.98 | 0.95 | 42 |

| species | precision | recall | f1-score | support |
|---|---|---|---|---|
| Campylorhamphus trochilirostris | 0.97 | 0.92 | 0.95 | 75 |
| Campylorhynchus griseus | 0.90 | 0.93 | 0.92 | 29 |
| Campylorhynchus turdinus | 0.97 | 0.95 | 0.96 | 75 |
| Campylorhynchus zonatus | 0.89 | 0.93 | 0.91 | 42 |
| Cantorchilus leucopogon | 0.97 | 0.91 | 0.94 | 35 |
| Cantorchilus leucotis | 0.92 | 0.99 | 0.95 | 75 |
| Cantorchilus nigricapillus | 0.95 | 0.92 | 0.93 | 75 |
| Capito auratus | 0.92 | 1.00 | 0.96 | 12 |
| Capsiempis flaveola | 0.94 | 0.89 | 0.92 | 75 |
| Caracara plancus | 0.95 | 0.93 | 0.94 | 45 |
| Catamblyrhynchus diadema | 0.94 | 1.00 | 0.97 | 29 |
| Catharus aurantiirostris | 0.93 | 0.95 | 0.94 | 75 |
| Catharus fuscater | 0.92 | 0.99 | 0.95 | 70 |
| Catharus fuscescens | 0.97 | 0.93 | 0.95 | 75 |
| Catharus minimus | 1.00 | 0.96 | 0.98 | 49 |
| Catharus ustulatus | 0.99 | 0.95 | 0.97 | 75 |
| Celeus flavus | 0.94 | 0.92 | 0.93 | 52 |
| Ceratopipra erythrocephala | 0.90 | 0.98 | 0.94 | 53 |
| Ceratopipra mentalis | 0.95 | 0.88 | 0.91 | 42 |
| Cercomacra cinerascens | 0.91 | 0.93 | 0.92 | 75 |
| Cercomacra nigricans | 0.88 | 0.94 | 0.91 | 32 |
| Cercomacroides fuscicauda | 0.94 | 0.98 | 0.96 | 46 |
| Cercomacroides nigrescens | 0.97 | 0.96 | 0.97 | 75 |
| Cercomacroides parkeri | 1.00 | 0.94 | 0.97 | 36 |
| Cercomacroides serva | 0.97 | 0.97 | 0.97 | 70 |
| Cercomacroides tyrannina | 0.93 | 0.88 | 0.90 | 75 |
| Certhiaxis cinnamomeus | 0.96 | 0.95 | 0.96 | 57 |
| Chamaeza campanisona | 0.97 | 0.95 | 0.96 | 75 |
| Chamaeza mollissima | 0.89 | 1.00 | 0.94 | 33 |
| Chamaeza nobilis | 1.00 | 1.00 | 1.00 | 21 |
| Chiroxiphia lanceolata | 0.98 | 0.96 | 0.97 | 52 |
| Chlorochrysa phoenicotis | 0.94 | 0.88 | 0.91 | 17 |
| Chlorophanes spiza | 0.92 | 0.96 | 0.94 | 25 |
| Chlorophonia cyanea | 1.00 | 0.95 | 0.97 | 19 |

| species | precision | recall | f1-score | support |
|---|---|---|---|---|
| Chlorophonia cyanocephala | 0.92 | 0.92 | 0.92 | 39 |
| Chlorophonia pyrrhophrys | 1.00 | 0.86 | 0.93 | 22 |
| Chlorornis riefferii | 0.94 | 1.00 | 0.97 | 29 |
| Chlorospingus canigularis | 0.96 | 0.88 | 0.92 | 25 |
| Chlorospingus flavigularis | 0.94 | 0.98 | 0.96 | 61 |
| Chlorospingus flavopectus | 0.96 | 0.91 | 0.93 | 75 |
| Chlorospingus semifuscus | 0.96 | 0.98 | 0.97 | 44 |
| Chrysomus icterocephalus | 0.89 | 0.86 | 0.88 | 29 |
| Cichlopsis leucogenys | 0.95 | 0.98 | 0.96 | 40 |
| Cinclodes excelsior | 0.92 | 0.92 | 0.92 | 13 |
| Cinnycerthia olivascens | 0.96 | 0.92 | 0.94 | 75 |
| Cinnycerthia unirufa | 0.95 | 0.97 | 0.96 | 39 |
| Cissopis leverianus | 1.00 | 0.80 | 0.89 | 30 |
| Cistothorus apolinari | 0.96 | 0.93 | 0.95 | 75 |
| Cistothorus platensis | 0.91 | 0.96 | 0.94 | 75 |
| Clibanornis rubiginosus | 0.96 | 0.99 | 0.97 | 75 |
| Clibanornis rufipectus | 1.00 | 0.89 | 0.94 | 18 |
| Clytoctantes alixii | 0.97 | 1.00 | 0.98 | 28 |
| Cnemotriccus fuscatus | 0.91 | 0.96 | 0.94 | 75 |
| Cnipodectes subbrunneus | 0.93 | 0.98 | 0.95 | 53 |
| Coereba flaveola | 0.97 | 0.89 | 0.93 | 75 |
| Colonia colonus | 0.81 | 0.93 | 0.87 | 28 |
| Conirostrum bicolor | 0.88 | 0.88 | 0.88 | 17 |
| Conirostrum speciosum | 0.98 | 0.94 | 0.96 | 50 |
| Conopias albovittatus | 0.81 | 0.93 | 0.87 | 14 |
| Conopias cinchoneti | 0.96 | 1.00 | 0.98 | 24 |
| Conopias parvus | 1.00 | 0.90 | 0.95 | 30 |
| Conopias trivirgatus | 0.93 | 1.00 | 0.97 | 14 |
| Conopophaga aurita | 1.00 | 0.90 | 0.95 | 20 |
| Conopophaga castaneiceps | 0.94 | 0.84 | 0.89 | 19 |
| Contopus cinereus | 1.00 | 0.87 | 0.93 | 31 |
| Contopus cooperi | 0.95 | 1.00 | 0.97 | 75 |
| Contopus fumigatus | 0.98 | 0.98 | 0.98 | 47 |
| Contopus sordidulus | 0.95 | 0.93 | 0.94 | 75 |

| species | precision | recall | f1-score | support |
|---|---|---|---|---|
| Contopus virens | 0.96 | 0.91 | 0.93 | 75 |
| Coryphospingus cucullatus | 0.99 | 0.96 | 0.97 | 69 |
| Coryphospingus pileatus | 0.98 | 0.98 | 0.98 | 42 |
| Corythopis torquatus | 0.98 | 0.98 | 0.98 | 60 |
| Cranioleuca curtata | 1.00 | 0.92 | 0.96 | 24 |
| Cranioleuca erythrops | 0.96 | 0.96 | 0.96 | 46 |
| Cranioleuca gutturata | 0.96 | 0.93 | 0.95 | 28 |
| Cranioleuca hellmayri | 1.00 | 0.94 | 0.97 | 31 |
| Cranioleuca vulpina | 0.88 | 0.98 | 0.93 | 59 |
| Cyanerpes caeruleus | 0.94 | 0.94 | 0.94 | 16 |
| Cyanerpes cyaneus | 0.82 | 1.00 | 0.90 | 23 |
| Cyanocorax affinis | 0.93 | 0.93 | 0.93 | 69 |
| Cyanocorax violaceus | 0.95 | 0.99 | 0.97 | 75 |
| Cyanocorax yncas | 0.96 | 0.95 | 0.95 | 75 |
| Cyanolyca armillata | 0.97 | 0.94 | 0.96 | 35 |
| Cyanolyca pulchra | 0.94 | 0.94 | 0.94 | 67 |
| Cyanolyca turcosa | 0.92 | 0.98 | 0.95 | 49 |
| Cyclarhis gujanensis | 0.91 | 0.91 | 0.91 | 75 |
| Cyclarhis nigrirostris | 0.92 | 0.94 | 0.93 | 62 |
| Cymbilaimus lineatus | 0.92 | 0.97 | 0.94 | 69 |
| Cyphorhinus arada | 0.97 | 0.96 | 0.97 | 75 |
| Cyphorhinus phaeocephalus | 0.99 | 0.97 | 0.98 | 75 |
| Cyphorhinus thoracicus | 0.96 | 0.97 | 0.97 | 75 |
| Dacnis cayana | 0.81 | 0.98 | 0.88 | 43 |
| Daptrius ater | 0.94 | 0.94 | 0.94 | 52 |
| Deconychura longicauda | 0.96 | 1.00 | 0.98 | 46 |
| Dendrexetastes rufigula | 0.52 | 0.98 | 0.68 | 49 |
| Dendrocincla fuliginosa | 0.90 | 0.87 | 0.88 | 75 |
| Dendrocincla tyrannina | 0.94 | 0.94 | 0.94 | 18 |
| Dendrocolaptes certhia | 0.99 | 0.91 | 0.94 | 75 |
| Dendrocolaptes picumnus | 0.97 | 0.97 | 0.97 | 37 |
| Dendrocolaptes sanctithomae | 0.97 | 0.94 | 0.96 | 35 |
| Dendroma rufa | 0.96 | 1.00 | 0.98 | 24 |
| Dendroplex picus | 0.93 | 0.88 | 0.90 | 75 |

| species | precision | recall | f1-score | support |
|---|---|---|---|---|
| Dichrozona cincta | 0.93 | 1.00 | 0.96 | 26 |
| Diglossa albilatera | 0.87 | 0.87 | 0.87 | 30 |
| Diglossa caerulescens | 1.00 | 0.79 | 0.88 | 28 |
| Diglossa cyanea | 0.87 | 0.86 | 0.86 | 56 |
| Diglossa glauca | 0.94 | 0.94 | 0.94 | 31 |
| Diglossa gloriosissima | 1.00 | 1.00 | 1.00 | 27 |
| Diglossa humeralis | 0.81 | 0.85 | 0.83 | 20 |
| Diglossa indigotica | 0.86 | 0.96 | 0.91 | 26 |
| Diglossa lafresnayii | 1.00 | 0.96 | 0.98 | 50 |
| Dives warczewiczi | 0.96 | 0.95 | 0.95 | 56 |
| Dolichonyx oryzivorus | 0.91 | 0.97 | 0.94 | 75 |
| Donacobius atricapilla | 0.90 | 0.97 | 0.94 | 75 |
| Drymophila devillei | 0.92 | 0.97 | 0.95 | 75 |
| Drymophila hellmayri | 0.92 | 0.90 | 0.91 | 40 |
| Drymophila striaticeps | 0.97 | 0.95 | 0.96 | 75 |
| Dubusia taeniata | 0.95 | 0.99 | 0.97 | 70 |
| Dumetella carolinensis | 0.96 | 0.95 | 0.95 | 75 |
| Dysithamnus leucostictus | 1.00 | 0.94 | 0.97 | 18 |
| Dysithamnus mentalis | 0.97 | 0.91 | 0.94 | 75 |
| Dysithamnus occidentalis | 1.00 | 0.97 | 0.98 | 30 |
| Dysithamnus puncticeps | 0.94 | 0.94 | 0.94 | 33 |
| Elaenia albiceps | 0.96 | 0.98 | 0.97 | 52 |
| Elaenia chiriquensis | 0.94 | 0.92 | 0.93 | 49 |
| Elaenia cristata | 0.96 | 0.96 | 0.96 | 27 |
| Elaenia flavogaster | 0.93 | 1.00 | 0.96 | 64 |
| Elaenia frantzii | 0.86 | 0.97 | 0.91 | 64 |
| Elaenia pallatangae | 1.00 | 0.98 | 0.99 | 41 |
| Elaenia parvirostris | 0.93 | 0.96 | 0.95 | 57 |
| Elaenia spectabilis | 0.97 | 1.00 | 0.99 | 35 |
| Emberizoides herbicola | 0.94 | 1.00 | 0.97 | 74 |
| Empidonax alnorum | 0.96 | 1.00 | 0.98 | 75 |
| Empidonax flaviventris | 0.95 | 1.00 | 0.98 | 62 |
| Empidonax minimus | 0.92 | 0.96 | 0.94 | 75 |
| Empidonax traillii | 0.95 | 0.95 | 0.95 | 75 |

| species | precision | recall | f1-score | support |
|---|---|---|---|---|
| Empidonax virescens | 0.97 | 0.96 | 0.97 | 75 |
| Empidonomus varius | 0.92 | 0.96 | 0.94 | 46 |
| Epinecrophylla fulviventris | 0.95 | 0.95 | 0.95 | 65 |
| Epinecrophylla haematonota | 1.00 | 1.00 | 1.00 | 19 |
| Epinecrophylla ornata | 1.00 | 1.00 | 1.00 | 46 |
| Epinecrophylla spodionota | 0.95 | 0.90 | 0.92 | 20 |
| Eremophila alpestris | 0.96 | 0.96 | 0.96 | 75 |
| Eucometis penicillata | 0.92 | 0.96 | 0.94 | 47 |
| Euphonia chlorotica | 0.94 | 0.89 | 0.92 | 75 |
| Euphonia chrysopasta | 0.96 | 0.96 | 0.96 | 56 |
| Euphonia concinna | 0.89 | 1.00 | 0.94 | 16 |
| Euphonia fulvicrissa | 0.93 | 0.93 | 0.93 | 15 |
| Euphonia laniirostris | 0.94 | 0.91 | 0.93 | 75 |
| Euphonia mesochrysa | 0.92 | 0.89 | 0.90 | 37 |
| Euphonia minuta | 0.94 | 0.86 | 0.90 | 37 |
| Euphonia plumbea | 1.00 | 1.00 | 1.00 | 27 |
| Euphonia rufiventris | 0.85 | 1.00 | 0.92 | 17 |
| Euphonia violacea | 0.90 | 0.85 | 0.88 | 75 |
| Euphonia xanthogaster | 0.99 | 0.93 | 0.96 | 75 |
| Euscarthmus meloryphus | 0.92 | 0.92 | 0.92 | 75 |
| Falco peregrinus | 1.00 | 0.96 | 0.98 | 75 |
| Fluvicola nengeta | 1.00 | 0.95 | 0.97 | 19 |
| Formicarius analis | 0.89 | 0.88 | 0.89 | 75 |
| Formicarius colma | 0.98 | 0.96 | 0.97 | 57 |
| Formicarius nigricapillus | 0.98 | 0.96 | 0.97 | 55 |
| Formicarius rufipectus | 0.91 | 0.95 | 0.93 | 61 |
| Formicivora grisea | 1.00 | 1.00 | 1.00 | 49 |
| Forpus conspicillatus | 0.95 | 0.90 | 0.93 | 21 |
| Frederickena fulva | 0.98 | 0.96 | 0.97 | 46 |
| Furnarius leucopus | 1.00 | 1.00 | 1.00 | 21 |
| Glyphorynchus spirurus | 0.92 | 0.97 | 0.94 | 68 |
| Grallaria alleni | 1.00 | 1.00 | 1.00 | 37 |
| Grallaria bangsi | 0.97 | 0.99 | 0.98 | 73 |
| Grallaria dignissima | 0.96 | 0.93 | 0.95 | 29 |

| species | precision | recall | f1-score | support |
|---|---|---|---|---|
| Grallaria flavotincta | 0.90 | 0.96 | 0.93 | 56 |
| Grallaria guatimalensis | 0.89 | 0.97 | 0.93 | 35 |
| Grallaria haplonota | 0.97 | 0.97 | 0.97 | 31 |
| Grallaria kaestneri | 0.96 | 1.00 | 0.98 | 26 |
| Grallaria milleri | 0.96 | 0.92 | 0.94 | 25 |
| Grallaria nuchalis | 0.92 | 0.95 | 0.93 | 75 |
| Grallaria quitensis | 0.91 | 0.95 | 0.93 | 75 |
| Grallaria ruficapilla | 0.94 | 0.88 | 0.91 | 75 |
| Grallaria rufula | 0.95 | 0.98 | 0.96 | 40 |
| Grallaria saturata | 0.98 | 0.98 | 0.98 | 43 |
| Grallaria squamigera | 0.92 | 1.00 | 0.96 | 48 |
| Grallaria varia | 0.98 | 0.96 | 0.97 | 47 |
| Grallaricula ferrugineipectus | 0.91 | 0.95 | 0.93 | 22 |
| Grallaricula flavirostris | 0.93 | 0.98 | 0.95 | 41 |
| Grallaricula lineifrons | 0.91 | 0.87 | 0.89 | 23 |
| Grallaricula nana | 0.95 | 0.95 | 0.95 | 75 |
| Gymnocichla nudiceps | 0.96 | 0.98 | 0.97 | 44 |
| Gymnopithys bicolor | 0.89 | 0.94 | 0.92 | 53 |
| Gymnopithys leucaspis | 1.00 | 0.90 | 0.95 | 20 |
| Gymnopithys rufigula | 0.93 | 0.87 | 0.90 | 31 |
| Haplospiza rustica | 0.94 | 0.94 | 0.94 | 50 |
| Hellmayrea gularis | 0.92 | 0.83 | 0.87 | 29 |
| Hemithraupis flavicollis | 0.89 | 0.84 | 0.86 | 19 |
| Hemithraupis guira | 0.89 | 0.92 | 0.91 | 53 |
| Hemitriccus granadensis | 0.97 | 0.95 | 0.96 | 37 |
| Hemitriccus iohannis | 0.96 | 0.92 | 0.94 | 24 |
| Hemitriccus margaritaceiventer | 0.95 | 0.95 | 0.95 | 75 |
| Hemitriccus striaticollis | 0.91 | 0.95 | 0.93 | 44 |
| Hemitriccus zosterops | 1.00 | 0.97 | 0.99 | 34 |
| Henicorhina anachoreta | 0.85 | 1.00 | 0.92 | 17 |
| Henicorhina leucophrys | 0.93 | 0.89 | 0.91 | 75 |
| Henicorhina leucosticta | 0.94 | 0.87 | 0.90 | 75 |
| Henicorhina negreti | 0.92 | 0.92 | 0.92 | 24 |
| Herpetotheres cachinnans | 0.96 | 1.00 | 0.98 | 75 |

| species | precision | recall | f1-score | support |
|---|---|---|---|---|
| Herpsilochmus rufimarginatus | 0.94 | 0.98 | 0.96 | 49 |
| Hirundinea ferruginea | 0.98 | 0.98 | 0.98 | 48 |
| Hirundo rustica | 0.92 | 0.97 | 0.95 | 75 |
| Hylocichla mustelina | 0.97 | 0.97 | 0.97 | 75 |
| Hylopezus fulviventris | 0.93 | 1.00 | 0.96 | 26 |
| Hylopezus macularius | 0.96 | 0.98 | 0.97 | 48 |
| Hylopezus perspicillatus | 0.90 | 1.00 | 0.95 | 75 |
| Hylophilus flavipes | 0.93 | 0.95 | 0.94 | 75 |
| Hylophilus olivaceus | 1.00 | 0.93 | 0.96 | 29 |
| Hylophilus semicinereus | 0.92 | 0.96 | 0.94 | 47 |
| Hylophilus thoracicus | 0.92 | 0.98 | 0.95 | 56 |
| Hylophylax naevioides | 0.96 | 0.88 | 0.92 | 60 |
| Hylophylax naevius | 0.95 | 0.95 | 0.95 | 75 |
| Hylophylax punctulatus | 0.97 | 0.96 | 0.97 | 75 |
| Hypocnemis flavescens | 1.00 | 0.95 | 0.97 | 20 |
| Hypocnemis hypoxantha | 0.96 | 1.00 | 0.98 | 55 |
| Hypocnemis peruviana | 0.93 | 0.93 | 0.93 | 75 |
| Hypocnemoides melanopogon | 1.00 | 0.90 | 0.95 | 40 |
| Ibycter americanus | 0.97 | 0.96 | 0.97 | 75 |
| Icterus cayanensis | 1.00 | 0.94 | 0.97 | 34 |
| Icterus chrysater | 0.98 | 0.92 | 0.95 | 50 |
| Icterus croconotus | 0.94 | 0.98 | 0.96 | 50 |
| Icterus galbula | 0.93 | 0.89 | 0.91 | 75 |
| Icterus mesomelas | 0.93 | 0.93 | 0.93 | 44 |
| Icterus nigrogularis | 0.92 | 0.97 | 0.95 | 37 |
| Icterus spurius | 0.93 | 0.89 | 0.91 | 75 |
| Inezia caudata | 1.00 | 1.00 | 1.00 | 14 |
| Inezia subflava | 0.88 | 0.78 | 0.82 | 9 |
| Iridosornis porphyrocephalus | 1.00 | 0.88 | 0.94 | 17 |
| Isleria guttata | 0.95 | 1.00 | 0.98 | 21 |
| Isleria hauxwelli | 0.95 | 0.94 | 0.95 | 65 |
| Ixothraupis rufigula | 0.88 | 0.82 | 0.85 | 17 |
| Kleinothraupis atropileus | 1.00 | 0.92 | 0.96 | 25 |
| Lamprospiza melanoleuca | 0.97 | 0.97 | 0.97 | 38 |

| species | precision | recall | f1-score | support |
|---|---|---|---|---|
| Lanio fulvus | 0.98 | 0.94 | 0.96 | 52 |
| Laniocera hypopyrra | 0.91 | 0.97 | 0.94 | 32 |
| Lathrotriccus euleri | 0.99 | 0.92 | 0.95 | 75 |
| Legatus leucophaius | 0.99 | 0.93 | 0.96 | 75 |
| Leistes bellicosus | 0.92 | 0.93 | 0.93 | 61 |
| Leistes militaris | 0.95 | 1.00 | 0.98 | 41 |
| Lepidocolaptes lacrymiger | 0.97 | 0.85 | 0.90 | 39 |
| Lepidocolaptes souleyetii | 0.95 | 0.90 | 0.92 | 40 |
| Lepidothrix coronata | 0.89 | 0.94 | 0.92 | 53 |
| Leptopogon amaurocephalus | 0.91 | 0.91 | 0.91 | 44 |
| Leptopogon rufipectus | 0.95 | 0.90 | 0.92 | 39 |
| Leptopogon superciliaris | 0.97 | 0.92 | 0.95 | 40 |
| Liosceles thoracicus | 0.96 | 0.97 | 0.97 | 75 |
| Lipaugus unirufus | 0.71 | 0.75 | 0.73 | 16 |
| Lipaugus vociferans | 0.94 | 0.96 | 0.95 | 75 |
| Lipaugus weberi | 1.00 | 1.00 | 1.00 | 12 |
| Lochmias nematura | 0.94 | 0.96 | 0.95 | 50 |
| Lophotriccus galeatus | 0.93 | 0.89 | 0.91 | 75 |
| Lophotriccus pileatus | 0.95 | 0.92 | 0.93 | 75 |
| Lophotriccus vitiosus | 0.94 | 0.97 | 0.96 | 33 |
| Loriotus cristatus | 0.88 | 0.88 | 0.88 | 33 |
| Loriotus luctuosus | 0.92 | 1.00 | 0.96 | 22 |
| Machaeropterus deliciosus | 1.00 | 0.96 | 0.98 | 48 |
| Machaeropterus regulus | 1.00 | 0.96 | 0.98 | 24 |
| Machetornis rixosa | 0.92 | 0.91 | 0.92 | 67 |
| Manacus manacus | 0.97 | 0.95 | 0.96 | 61 |
| Mazaria propinqua | 0.91 | 0.91 | 0.91 | 22 |
| Mecocerculus leucophrys | 0.96 | 0.95 | 0.95 | 75 |
| Mecocerculus stictopterus | 0.80 | 0.92 | 0.85 | 38 |
| Megarynchus pitangua | 0.95 | 0.97 | 0.96 | 75 |
| Melanerpes formicivorus | 0.94 | 0.99 | 0.96 | 75 |
| Melanopareia elegans | 0.92 | 0.96 | 0.94 | 50 |
| Melanospiza bicolor | 0.70 | 0.78 | 0.74 | 9 |
| Micrastur ruficollis | 0.99 | 1.00 | 0.99 | 75 |

| species | precision | recall | f1-score | support |
|---|---|---|---|---|
| Microbates cinereiventris | 0.98 | 0.96 | 0.97 | 49 |
| Microbates collaris | 0.94 | 0.97 | 0.95 | 30 |
| Microcerculus bambla | 0.94 | 0.96 | 0.95 | 70 |
| Microcerculus marginatus | 0.95 | 0.93 | 0.94 | 75 |
| Microrhopias quixensis | 0.92 | 0.92 | 0.92 | 75 |
| Milvago chimachima | 0.93 | 0.89 | 0.91 | 75 |
| Mimus gilvus | 0.92 | 0.93 | 0.93 | 75 |
| Mionectes oleagineus | 0.95 | 0.98 | 0.97 | 60 |
| Mionectes striaticollis | 0.89 | 0.96 | 0.92 | 25 |
| Mitrephanes phaeocercus | 0.90 | 1.00 | 0.95 | 37 |
| Molothrus bonariensis | 0.90 | 0.99 | 0.94 | 75 |
| Molothrus oryzivorus | 0.88 | 0.70 | 0.78 | 20 |
| Myadestes ralloides | 0.88 | 0.89 | 0.89 | 75 |
| Myiarchus cephalotes | 1.00 | 0.96 | 0.98 | 24 |
| Myiarchus crinitus | 0.95 | 0.97 | 0.96 | 75 |
| Myiarchus ferox | 0.96 | 0.93 | 0.95 | 75 |
| Myiarchus tuberculifer | 0.90 | 0.99 | 0.94 | 75 |
| Myiarchus tyrannulus | 0.96 | 0.95 | 0.95 | 75 |
| Myiodynastes chrysocephalus | 0.95 | 1.00 | 0.97 | 75 |
| Myiodynastes luteiventris | 0.95 | 0.96 | 0.95 | 75 |
| Myiodynastes maculatus | 0.92 | 0.92 | 0.92 | 75 |
| Myiopagis caniceps | 0.92 | 0.84 | 0.88 | 57 |
| Myiopagis flavivertex | 0.92 | 1.00 | 0.96 | 11 |
| Myiopagis gaimardii | 0.91 | 0.94 | 0.92 | 64 |
| Myiopagis subplacens | 0.88 | 0.88 | 0.88 | 25 |
| Myiopagis viridicata | 0.95 | 0.96 | 0.95 | 75 |
| Myiophobus cryptoxanthus | 0.95 | 0.72 | 0.82 | 25 |
| Myiophobus fasciatus | 0.97 | 0.95 | 0.96 | 73 |
| Myiophobus flavicans | 0.97 | 0.89 | 0.93 | 36 |
| Myiophobus phoenicomitra | 0.93 | 1.00 | 0.97 | 28 |
| Myiophobus roraimae | 0.82 | 0.96 | 0.88 | 24 |
| Myiornis atricapillus | 0.97 | 1.00 | 0.99 | 37 |
| Myiornis ecaudatus | 1.00 | 0.95 | 0.97 | 38 |
| Myiotheretes fumigatus | 0.96 | 1.00 | 0.98 | 23 |

| species | precision | recall | f1-score | support |
|---|---|---|---|---|
| Myiotheretes striaticollis | 1.00 | 0.90 | 0.95 | 20 |
| Myiotriccus ornatus | 0.95 | 0.98 | 0.97 | 61 |
| Myiozetetes cayanensis | 0.99 | 0.93 | 0.96 | 75 |
| Myiozetetes luteiventris | 1.00 | 1.00 | 1.00 | 24 |
| Myiozetetes similis | 0.97 | 0.92 | 0.95 | 75 |
| Myornis senilis | 0.94 | 0.94 | 0.94 | 51 |
| Myrmeciza longipes | 0.99 | 0.95 | 0.97 | 75 |
| Myrmelastes hyperythrus | 0.95 | 0.96 | 0.95 | 75 |
| Myrmelastes leucostigma | 0.95 | 0.96 | 0.95 | 75 |
| Myrmelastes schistaceus | 1.00 | 1.00 | 1.00 | 21 |
| Myrmoborus leucophrys | 0.92 | 0.93 | 0.93 | 75 |
| Myrmoborus lugubris | 0.94 | 0.97 | 0.96 | 33 |
| Myrmoborus myotherinus | 0.97 | 0.93 | 0.95 | 75 |
| Myrmochanes hemileucus | 1.00 | 0.94 | 0.97 | 17 |
| Myrmophylax atrothorax | 0.95 | 0.93 | 0.94 | 75 |
| Myrmornis torquata | 0.97 | 1.00 | 0.98 | 29 |
| Myrmothera campanisona | 0.93 | 0.95 | 0.94 | 75 |
| Myrmotherula assimilis | 0.91 | 0.97 | 0.94 | 30 |
| Myrmotherula axillaris | 0.95 | 0.93 | 0.94 | 75 |
| Myrmotherula brachyura | 0.93 | 0.98 | 0.96 | 56 |
| Myrmotherula ignota | 0.97 | 0.94 | 0.95 | 33 |
| Myrmotherula longicauda | 1.00 | 1.00 | 1.00 | 20 |
| Myrmotherula longipennis | 0.96 | 0.96 | 0.96 | 23 |
| Myrmotherula menetriesii | 0.96 | 0.94 | 0.95 | 52 |
| Myrmotherula multostriata | 0.96 | 0.95 | 0.95 | 56 |
| Myrmotherula pacifica | 0.95 | 1.00 | 0.98 | 20 |
| Myrmotherula schisticolor | 0.96 | 0.92 | 0.94 | 24 |
| Nasica longirostris | 0.92 | 0.87 | 0.89 | 39 |
| Nemosia pileata | 0.84 | 0.95 | 0.89 | 22 |
| Neoctantes niger | 0.93 | 1.00 | 0.96 | 26 |
| Neopelma chrysocephalum | 0.88 | 0.98 | 0.92 | 44 |
| Neopipo cinnamomea | 0.97 | 1.00 | 0.98 | 28 |
| Nephelomyias pulcher | 1.00 | 0.86 | 0.92 | 14 |
| Ochthoeca cinnamomeiventris | 1.00 | 0.96 | 0.98 | 28 |

| species | precision | recall | f1-score | support |
|---|---|---|---|---|
| Ochthoeca fumicolor | 1.00 | 0.93 | 0.96 | 14 |
| Odontorchilus branickii | 0.89 | 0.94 | 0.91 | 17 |
| Oncostoma cinereigulare | 1.00 | 0.86 | 0.92 | 21 |
| Oreothraupis arremonops | 1.00 | 0.97 | 0.98 | 33 |
| Ornithion inerme | 0.95 | 0.97 | 0.96 | 65 |
| Pachyramphus albogriseus | 0.83 | 1.00 | 0.91 | 20 |
| Pachyramphus cinnamomeus | 0.93 | 0.93 | 0.93 | 14 |
| Pachyramphus minor | 0.93 | 0.82 | 0.88 | 17 |
| Pachyramphus polychopterus | 0.91 | 0.89 | 0.90 | 44 |
| Pachysylvia decurtata | 0.95 | 1.00 | 0.97 | 37 |
| Pachysylvia hypoxantha | 0.87 | 0.95 | 0.91 | 21 |
| Pachysylvia muscicapina | 1.00 | 0.88 | 0.94 | 17 |
| Passer domesticus | 0.96 | 0.97 | 0.97 | 75 |
| Passerculus sandwichensis | 0.96 | 0.96 | 0.96 | 75 |
| Percnostola rufifrons | 0.99 | 0.95 | 0.97 | 75 |
| Perissocephalus tricolor | 1.00 | 1.00 | 1.00 | 27 |
| Petrochelidon pyrrhonota | 0.96 | 0.97 | 0.97 | 75 |
| Phacellodomus rufifrons | 0.91 | 0.98 | 0.94 | 40 |
| Phaenostictus mcleannani | 0.97 | 1.00 | 0.98 | 32 |
| Phaeomyias murina | 0.93 | 0.87 | 0.90 | 75 |
| Pheugopedius coraya | 0.96 | 0.92 | 0.94 | 75 |
| Pheugopedius euophrys | 0.99 | 0.96 | 0.97 | 75 |
| Pheugopedius fasciatoventris | 0.95 | 0.95 | 0.95 | 75 |
| Pheugopedius genibarbis | 0.95 | 0.95 | 0.95 | 75 |
| Pheugopedius mystacalis | 0.92 | 0.93 | 0.93 | 75 |
| Pheugopedius rutilus | 0.93 | 0.91 | 0.92 | 75 |
| Pheugopedius sclateri | 0.99 | 0.88 | 0.93 | 75 |
| Pheugopedius spadix | 0.93 | 0.97 | 0.95 | 29 |
| Philydor erythrocercum | 0.95 | 1.00 | 0.97 | 18 |
| Phlegopsis nigromaculata | 0.98 | 0.93 | 0.95 | 55 |
| Phoenicircus nigricollis | 0.93 | 0.93 | 0.93 | 14 |
| Phyllomyias burmeisteri | 0.95 | 0.95 | 0.95 | 42 |
| Phyllomyias griseiceps | 0.96 | 0.96 | 0.96 | 45 |
| Phyllomyias nigrocapillus | 0.96 | 0.88 | 0.92 | 26 |

| species | precision | recall | f1-score | support |
|---|---|---|---|---|
| Phylloscartes gualaquizae | 0.89 | 1.00 | 0.94 | 16 |
| Piculus flavigula | 0.83 | 0.77 | 0.80 | 13 |
| Picumnus exilis | 0.92 | 0.92 | 0.92 | 25 |
| Pionus menstruus | 0.92 | 0.97 | 0.95 | 75 |
| Pipraeidea melanonota | 0.81 | 0.90 | 0.85 | 39 |
| Pipreola arcuata | 0.94 | 0.97 | 0.95 | 31 |
| Pipreola riefferii | 0.83 | 0.96 | 0.89 | 26 |
| Piprites chloris | 0.95 | 0.95 | 0.95 | 20 |
| Pitangus sulphuratus | 0.96 | 0.97 | 0.97 | 75 |
| Pittasoma rufopileatum | 0.93 | 1.00 | 0.96 | 26 |
| Platyrinchus coronatus | 0.96 | 0.96 | 0.96 | 24 |
| Platyrinchus mystaceus | 0.95 | 0.96 | 0.95 | 54 |
| Platyrinchus platyrhynchos | 0.95 | 1.00 | 0.98 | 42 |
| Ploceus cucullatus | 0.96 | 0.95 | 0.95 | 75 |
| Poecilotriccus capitalis | 1.00 | 0.94 | 0.97 | 17 |
| Poecilotriccus ruficeps | 0.87 | 0.93 | 0.90 | 14 |
| Poecilotriccus sylvia | 1.00 | 0.83 | 0.91 | 12 |
| Poliocrania exsul | 0.90 | 0.87 | 0.88 | 75 |
| Polioptila plumbea | 0.94 | 0.87 | 0.90 | 75 |
| Premnoplex brunnescens | 0.90 | 0.96 | 0.93 | 48 |
| Procnias averano | 0.89 | 1.00 | 0.94 | 51 |
| Progne chalybea | 0.92 | 0.91 | 0.92 | 54 |
| Progne subis | 0.95 | 0.97 | 0.96 | 75 |
| Progne tapera | 0.92 | 0.90 | 0.91 | 49 |
| Psarocolius angustifrons | 0.95 | 0.84 | 0.89 | 75 |
| Psarocolius bifasciatus | 0.97 | 0.94 | 0.95 | 31 |
| Psarocolius decumanus | 0.97 | 0.97 | 0.97 | 75 |
| Psarocolius viridis | 0.88 | 0.90 | 0.89 | 51 |
| Psarocolius wagleri | 0.93 | 0.98 | 0.95 | 51 |
| Pseudocolaptes boissonneautii | 1.00 | 0.95 | 0.97 | 19 |
| Pseudopipra pipra | 0.93 | 0.98 | 0.95 | 41 |
| Pseudotriccus pelzelni | 0.96 | 0.94 | 0.95 | 47 |
| Pseudotriccus ruficeps | 0.94 | 0.92 | 0.93 | 48 |
| Pteroglossus castanotis | 0.98 | 1.00 | 0.99 | 53 |

| species | precision | recall | f1-score | support |
|---|---|---|---|---|
| Pygiptila stellaris | 1.00 | 0.90 | 0.95 | 39 |
| Pygochelidon cyanoleuca | 0.98 | 0.96 | 0.97 | 55 |
| Pyriglena leuconota | 0.94 | 0.96 | 0.95 | 75 |
| Pyriglena maura | 0.97 | 0.95 | 0.96 | 75 |
| Pyrrhomyias cinnamomeus | 0.88 | 0.92 | 0.90 | 75 |
| Querula purpurata | 0.95 | 0.95 | 0.95 | 61 |
| Quiscalus lugubris | 1.00 | 0.92 | 0.96 | 13 |
| Quiscalus mexicanus | 0.95 | 0.92 | 0.93 | 75 |
| Ramphastos tucanus | 0.99 | 0.99 | 0.99 | 75 |
| Ramphocaenus melanurus | 0.94 | 0.91 | 0.93 | 75 |
| Ramphocelus carbo | 0.86 | 0.93 | 0.90 | 75 |
| Ramphocelus dimidiatus | 1.00 | 0.83 | 0.90 | 23 |
| Ramphotrigon fuscicauda | 0.97 | 0.94 | 0.95 | 33 |
| Ramphotrigon megacephalum | 0.97 | 0.92 | 0.95 | 39 |
| Ramphotrigon ruficauda | 0.98 | 1.00 | 0.99 | 65 |
| Rhegmatorhina melanosticta | 0.92 | 0.92 | 0.92 | 26 |
| Rhodinocichla rosea | 0.97 | 0.95 | 0.96 | 75 |
| Rhynchocyclus olivaceus | 0.97 | 0.91 | 0.94 | 33 |
| Rhytipterna simplex | 0.96 | 0.94 | 0.95 | 52 |
| Riparia riparia | 0.95 | 0.96 | 0.95 | 75 |
| Rupicola peruvianus | 0.96 | 0.95 | 0.95 | 75 |
| Rupicola rupicola | 0.95 | 1.00 | 0.97 | 35 |
| Sakesphorus canadensis | 0.98 | 0.95 | 0.97 | 60 |
| Saltator atripennis | 0.92 | 1.00 | 0.96 | 48 |
| Saltator coerulescens | 0.95 | 0.93 | 0.94 | 75 |
| Saltator grossus | 0.90 | 0.92 | 0.91 | 75 |
| Saltator maximus | 0.89 | 0.96 | 0.92 | 75 |
| Saltator striatipectus | 0.92 | 0.95 | 0.93 | 75 |
| Sayornis nigricans | 0.97 | 0.99 | 0.98 | 75 |
| Schiffornis aenea | 0.90 | 0.93 | 0.92 | 29 |
| Schiffornis major | 1.00 | 0.79 | 0.88 | 33 |
| Schiffornis turdina | 0.94 | 0.97 | 0.95 | 75 |
| Schiffornis veraepacis | 0.96 | 0.93 | 0.95 | 57 |
| Schistochlamys melanopis | 1.00 | 0.86 | 0.93 | 29 |

| species | precision | recall | f1-score | support |
|---|---|---|---|---|
| Sciaphylax castanea | 0.98 | 0.92 | 0.95 | 51 |
| Sclateria naevia | 0.94 | 0.96 | 0.95 | 75 |
| Sclerurus albigularis | 1.00 | 0.95 | 0.97 | 73 |
| Sclerurus caudacutus | 0.95 | 0.98 | 0.96 | 40 |
| Sclerurus guatemalensis | 1.00 | 1.00 | 1.00 | 32 |
| Sclerurus mexicanus | 1.00 | 0.90 | 0.95 | 20 |
| Sclerurus obscurior | 0.94 | 0.96 | 0.95 | 75 |
| Sclerurus rufigularis | 0.97 | 0.90 | 0.93 | 39 |
| Scytalopus alvarezlopezi | 1.00 | 1.00 | 1.00 | 35 |
| Scytalopus atratus | 0.96 | 1.00 | 0.98 | 75 |
| Scytalopus chocoensis | 1.00 | 0.94 | 0.97 | 34 |
| Scytalopus griseicollis | 0.98 | 0.90 | 0.94 | 50 |
| Scytalopus latebricola | 1.00 | 0.94 | 0.97 | 48 |
| Scytalopus latrans | 0.96 | 0.99 | 0.97 | 75 |
| Scytalopus micropterus | 0.95 | 0.95 | 0.95 | 55 |
| Scytalopus opacus | 0.92 | 0.92 | 0.92 | 24 |
| Scytalopus perijanus | 1.00 | 1.00 | 1.00 | 16 |
| Scytalopus rodriguezi | 1.00 | 1.00 | 1.00 | 30 |
| Scytalopus sanctaemartae | 0.95 | 0.95 | 0.95 | 22 |
| Scytalopus spillmanni | 0.96 | 0.93 | 0.95 | 75 |
| Scytalopus stilesi | 1.00 | 0.95 | 0.98 | 22 |
| Scytalopus vicinior | 0.97 | 0.99 | 0.98 | 72 |
| Selenidera reinwardtii | 0.92 | 0.92 | 0.92 | 24 |
| Sericossypha albocristata | 0.99 | 0.99 | 0.99 | 75 |
| Sicalis citrina | 0.95 | 0.93 | 0.94 | 41 |
| Sicalis flaveola | 0.97 | 0.96 | 0.97 | 75 |
| Sicalis luteola | 0.96 | 0.96 | 0.96 | 56 |
| Sipia berlepschi | 1.00 | 0.98 | 0.99 | 43 |
| Sipia laemosticta | 0.96 | 0.94 | 0.95 | 54 |
| Sipia nigricauda | 0.98 | 0.94 | 0.96 | 47 |
| Sipia palliata | 0.95 | 1.00 | 0.98 | 40 |
| Sittasomus griseicapillus | 0.94 | 0.89 | 0.92 | 75 |
| Snowornis subalaris | 1.00 | 0.88 | 0.93 | 8 |
| Sphenopsis frontalis | 1.00 | 0.95 | 0.98 | 22 |

| species | precision | recall | f1-score | support |
|---|---|---|---|---|
| Sphenopsis melanotis | 1.00 | 1.00 | 1.00 | 26 |
| Spinus magellanicus | 0.90 | 0.94 | 0.92 | 48 |
| Spinus olivaceus | 1.00 | 1.00 | 1.00 | 17 |
| Spinus psaltria | 0.96 | 0.92 | 0.94 | 75 |
| Spizella pallida | 0.94 | 0.96 | 0.95 | 75 |
| Sporathraupis cyanocephala | 0.89 | 1.00 | 0.94 | 17 |
| Sporophila angolensis | 0.93 | 0.95 | 0.94 | 75 |
| Sporophila caerulescens | 0.88 | 0.95 | 0.91 | 60 |
| Sporophila castaneiventris | 0.94 | 0.89 | 0.92 | 19 |
| Sporophila corvina | 0.89 | 0.86 | 0.88 | 29 |
| Sporophila funerea | 0.97 | 0.94 | 0.95 | 31 |
| Sporophila lineola | 0.95 | 0.95 | 0.95 | 63 |
| Sporophila luctuosa | 0.88 | 1.00 | 0.94 | 38 |
| Sporophila minuta | 0.93 | 0.94 | 0.93 | 53 |
| Sporophila nigricollis | 0.95 | 0.93 | 0.94 | 75 |
| Sporophila plumbea | 0.93 | 0.95 | 0.94 | 40 |
| Sporophila schistacea | 0.97 | 0.93 | 0.95 | 41 |
| Sporophila telasco | 1.00 | 0.91 | 0.95 | 22 |
| Stelgidopteryx ruficollis | 0.98 | 0.93 | 0.95 | 44 |
| Stelgidopteryx serripennis | 1.00 | 0.96 | 0.98 | 24 |
| Stigmatura napensis | 1.00 | 1.00 | 1.00 | 14 |
| Stilpnia cayana | 0.87 | 0.87 | 0.87 | 39 |
| Stilpnia cyanicollis | 0.95 | 0.95 | 0.95 | 21 |
| Stilpnia heinei | 0.95 | 0.95 | 0.95 | 20 |
| Stilpnia larvata | 0.81 | 0.89 | 0.85 | 19 |
| Stilpnia vitriolina | 0.90 | 0.82 | 0.86 | 11 |
| Sturnella magna | 0.92 | 0.91 | 0.91 | 75 |
| Sublegatus modestus | 0.96 | 0.88 | 0.92 | 26 |
| Synallaxis albescens | 0.96 | 0.91 | 0.93 | 75 |
| Synallaxis albigularis | 1.00 | 1.00 | 1.00 | 70 |
| Synallaxis azarae | 0.97 | 0.96 | 0.97 | 75 |
| Synallaxis brachyura | 0.96 | 0.96 | 0.96 | 52 |
| Synallaxis candei | 1.00 | 0.88 | 0.94 | 17 |
| Synallaxis cherriei | 1.00 | 0.98 | 0.99 | 40 |

| species | precision | recall | f1-score | support |
|---|---|---|---|---|
| Synallaxis cinnamomea | 0.95 | 0.97 | 0.96 | 38 |
| Synallaxis fuscorufa | 1.00 | 0.97 | 0.99 | 36 |
| Synallaxis gujanensis | 0.97 | 0.97 | 0.97 | 70 |
| Synallaxis rutilans | 0.98 | 0.96 | 0.97 | 51 |
| Synallaxis unirufa | 0.92 | 0.97 | 0.94 | 68 |
| Syndactyla subalaris | 0.94 | 0.85 | 0.89 | 40 |
| Tachycineta albilinea | 0.82 | 0.93 | 0.88 | 15 |
| Tachycineta albiventer | 1.00 | 0.93 | 0.97 | 15 |
| Tachycineta bicolor | 0.95 | 0.95 | 0.95 | 75 |
| Tachycineta thalassina | 0.94 | 0.98 | 0.96 | 65 |
| Tachyphonus delatrii | 0.92 | 1.00 | 0.96 | 35 |
| Tachyphonus rufus | 0.93 | 0.93 | 0.93 | 41 |
| Tangara arthus | 0.94 | 0.83 | 0.88 | 18 |
| Tangara chilensis | 0.94 | 0.98 | 0.96 | 47 |
| Tangara gyrola | 0.95 | 0.92 | 0.93 | 38 |
| Tangara labradorides | 0.88 | 0.83 | 0.86 | 18 |
| Tangara nigroviridis | 1.00 | 0.87 | 0.93 | 15 |
| Tangara parzudakii | 0.87 | 0.81 | 0.84 | 16 |
| Tangara velia | 0.96 | 1.00 | 0.98 | 22 |
| Taraba major | 0.93 | 0.92 | 0.93 | 75 |
| Terenotriccus erythrurus | 0.93 | 1.00 | 0.97 | 14 |
| Tersina viridis | 0.95 | 0.91 | 0.93 | 45 |
| Thamnistes anabatinus | 0.89 | 0.93 | 0.91 | 27 |
| Thamnomanes ardesiacus | 0.97 | 0.94 | 0.95 | 62 |
| Thamnomanes caesius | 0.96 | 0.95 | 0.95 | 74 |
| Thamnophilus aethiops | 0.95 | 0.97 | 0.96 | 61 |
| Thamnophilus amazonicus | 0.88 | 0.97 | 0.92 | 37 |
| Thamnophilus atrinucha | 0.96 | 0.94 | 0.95 | 71 |
| Thamnophilus doliatus | 0.97 | 0.91 | 0.94 | 75 |
| Thamnophilus multistriatus | 0.94 | 0.98 | 0.96 | 51 |
| Thamnophilus murinus | 0.98 | 1.00 | 0.99 | 46 |
| Thamnophilus nigrocinereus | 0.96 | 0.89 | 0.93 | 28 |
| Thamnophilus punctatus | 0.96 | 0.94 | 0.95 | 54 |
| Thamnophilus schistaceus | 1.00 | 0.91 | 0.95 | 44 |

| species | precision | recall | f1-score | support |
|---|---|---|---|---|
| Thamnophilus tenuepunctatus | 0.97 | 0.97 | 0.97 | 33 |
| Thamnophilus unicolor | 0.96 | 0.90 | 0.93 | 30 |
| Thlypopsis sordida | 0.97 | 0.86 | 0.91 | 37 |
| Thlypopsis superciliaris | 0.97 | 0.94 | 0.95 | 33 |
| Thraupis episcopus | 0.92 | 0.88 | 0.90 | 75 |
| Thraupis palmarum | 0.93 | 0.89 | 0.91 | 75 |
| Thripadectes flammulatus | 0.94 | 0.97 | 0.96 | 34 |
| Thripadectes holostictus | 0.90 | 0.96 | 0.93 | 75 |
| Thripadectes ignobilis | 0.96 | 0.85 | 0.90 | 27 |
| Thripadectes melanorhynchus | 1.00 | 0.96 | 0.98 | 27 |
| Thripadectes virgaticeps | 0.96 | 0.98 | 0.97 | 48 |
| Thripophaga fusciceps | 0.96 | 0.90 | 0.93 | 29 |
| Thryophilus nicefori | 0.90 | 0.95 | 0.92 | 19 |
| Thryophilus rufalbus | 0.97 | 0.96 | 0.97 | 75 |
| Tiaris olivaceus | 1.00 | 0.93 | 0.97 | 45 |
| Tityra semifasciata | 0.94 | 0.92 | 0.93 | 36 |
| Todirostrum chrysocrotaphum | 0.91 | 0.97 | 0.94 | 31 |
| Todirostrum cinereum | 0.94 | 0.85 | 0.90 | 75 |
| Todirostrum maculatum | 1.00 | 0.94 | 0.97 | 16 |
| Todirostrum nigriceps | 0.89 | 0.94 | 0.92 | 18 |
| Todirostrum pictum | 0.94 | 1.00 | 0.97 | 16 |
| Tolmomyias assimilis | 0.93 | 0.89 | 0.91 | 46 |
| Tolmomyias flaviventris | 0.89 | 1.00 | 0.94 | 51 |
| Tolmomyias poliocephalus | 0.95 | 0.93 | 0.94 | 45 |
| Tolmomyias sulphurescens | 0.92 | 0.92 | 0.92 | 75 |
| Troglodytes aedon | 0.97 | 0.91 | 0.94 | 75 |
| Troglodytes solstitialis | 0.95 | 0.97 | 0.96 | 63 |
| Tunchiornis ochraceiceps | 0.94 | 0.97 | 0.96 | 66 |
| Tyranneutes stolzmanni | 0.97 | 0.95 | 0.96 | 66 |
| Tyrannopsis sulphurea | 0.93 | 0.95 | 0.94 | 40 |
| Tyrannus dominicensis | 0.97 | 1.00 | 0.98 | 63 |
| Tyrannus forficatus | 1.00 | 0.88 | 0.94 | 25 |
| Tyrannus melancholicus | 0.99 | 0.93 | 0.96 | 75 |
| Tyrannus niveigularis | 0.86 | 0.95 | 0.90 | 19 |

| species | precision | recall | f1-score | support |
|---|---|---|---|---|
| Tyrannus savana | 0.89 | 0.94 | 0.91 | 33 |
| Tyrannus tyrannus | 0.98 | 0.96 | 0.97 | 53 |
| Vireo altiloquus | 0.96 | 0.93 | 0.94 | 55 |
| Vireo chivi | 0.96 | 0.95 | 0.95 | 75 |
| Vireo flavifrons | 0.96 | 0.95 | 0.95 | 75 |
| Vireo flavoviridis | 0.99 | 0.93 | 0.96 | 75 |
| Vireo gilvus | 0.94 | 0.97 | 0.95 | 75 |
| Vireo griseus | 0.97 | 0.92 | 0.95 | 75 |
| Vireo leucophrys | 0.98 | 0.94 | 0.96 | 52 |
| Vireo masteri | 0.97 | 1.00 | 0.99 | 33 |
| Vireo olivaceus | 0.99 | 0.93 | 0.96 | 75 |
| Vireo philadelphicus | 0.94 | 0.97 | 0.96 | 67 |
| Vireolanius leucotis | 0.95 | 0.98 | 0.96 | 56 |
| Volatinia jacarina | 0.96 | 0.93 | 0.95 | 75 |
| Willisornis poecilinotus | 0.96 | 0.97 | 0.97 | 73 |
| Xenops minutus | 0.96 | 0.92 | 0.94 | 75 |
| Xenops rutilans | 0.87 | 0.96 | 0.91 | 75 |
| Xiphocolaptes promeropirhynchus | 0.97 | 0.92 | 0.95 | 75 |
| Xiphorhynchus elegans | 1.00 | 0.90 | 0.95 | 21 |
| Xiphorhynchus erythropygius | 0.97 | 0.97 | 0.97 | 34 |
| Xiphorhynchus guttatus | 0.93 | 0.89 | 0.91 | 75 |
| Xiphorhynchus lachrymosus | 1.00 | 0.89 | 0.94 | 18 |
| Xiphorhynchus obsoletus | 0.96 | 0.93 | 0.94 | 27 |
| Xiphorhynchus pardalotus | 1.00 | 0.88 | 0.94 | 17 |
| Xiphorhynchus susurrans | 0.90 | 0.95 | 0.92 | 75 |
| Xiphorhynchus triangularis | 0.90 | 0.96 | 0.93 | 28 |
| Zimmerius chrysops | 0.89 | 0.93 | 0.91 | 45 |
| Zonotrichia capensis | 0.92 | 0.95 | 0.93 | 75 |
| Zonotrichia leucophrys | 0.93 | 0.95 | 0.94 | 75 |
